# Circular RNA profiling identifies *circ5078* as a *BMPR2*-derived regulator of endothelial proliferation and stress responses

**DOI:** 10.1101/2023.09.18.556866

**Authors:** M. Martin VandenBroek, Mackenzie C. Sharp, Patrick Thompson, Emmanuel Fagbola, Douglas Quilty, Jeffrey D. Mewburn, Anne L. Theilmann, Kimberly Dunham-Snary, Stephen L. Archer, Neil Renwick, Mark L. Ormiston

## Abstract

Germline loss-of-function *BMPR2* mutations are the leading genetic cause of pulmonary arterial hypertension (PAH) and are strongly linked to aberrant endothelial proliferation and impaired translational stress responses. While these effects are generally attributed to a loss of the type-II bone morphogenetic protein receptor (BMPR-II), we used circular RNA profiling to identify *circ5078* as a new functional RNA derived from exon 12 of the *BMPR2* gene. *circ5078* and linear *BMPR2* mRNA exert opposing effects on endothelial proliferation and stress granule formation, driving impaired stress responses in PAH patient-derived endothelial cells that are deficient in linear *BMPR2* transcripts, but not *circ5078*. Rebalancing circular to linear transcript abundance by *circ5078* depletion rescued stress granule formation in patient-derived endothelial cells, independent of BMPR-II protein levels. Polysome analysis demonstrated a reduction in free ribosomal subunits with the depletion of either linear or circular *BMPR2* transcripts, and an accumulation of 80S monosomes exclusively with linear *BMPR2* mRNA loss. These effects did not impact global protein synthesis or stress-induced eIF2α phosphorylation, but did alter the translational efficiency of multiple genes, including a group of nuclear encoded, mitochondrial ribosome proteins that were translationally enhanced with *circ5078* silencing. Endothelial *circ5078* depletion increased the efficiency of oxidative metabolism while reducing mitochondrial spare capacity, providing a potential link between mitochondrial function, proliferation and translational stress responses. Together, these findings reveal interdependent roles for linear and circular *BMPR2* transcripts as functional contributors to the endothelial phenotype of PAH.

## Introduction

Mutations in *BMPR2*, the gene encoding the bone morphogenetic protein (BMP) type-II receptor (BMPR-II), are strongly linked to pulmonary arterial hypertension (PAH), a disease of occlusive pulmonary vascular remodelling that is marked by the obstructive proliferation of endothelial and vascular smooth muscle cells. Heterozygous loss of function *BMPR2* mutations, including deletions, insertions and point-mutations across all exons and many noncoding regions of the gene^1,2^, account for 70% of the heritable form of PAH, and 20% of seemingly idiopathic disease^3^. While the excessive muscularization of distal pulmonary arteries is a prominent feature of established PAH, genetic and experimental evidence point to *BMPR2* loss in the pulmonary endothelium as an essential contributor to vascular inflammation and pathological remodeling^4^.

In addition to aberrant proliferation and an enhanced susceptibility to apoptosis, endothelial *BMPR2* loss is also linked to an impairment of stress-induced translational repression that is characterized by reductions in both stress granule formation and eIF2α phosphorylation^5–10^. Biallelic mutations in *EIF2AK4*, the gene encoding the eIF2α kinase, GCN2, have also been identified in patients with pulmonary veno-occlusive disease (PVOD), a rare subtype of PAH, alongside compound heterozygous *EIF2AK4*/*BMPR2* mutations in a small group of PAH patients^11^. Although these findings highlight a connection between the genetic drivers of pulmonary vascular disease and altered translational control, the precise mechanisms by which *BMPR2* mutations influence endothelial translation and stress responses remain uncertain. To date, investigations into these mechanisms have primarily focused on a loss of canonical and non-canonical signalling via the BMPR-II receptor^12–14^, which can be encoded by a full-length transcript (*BMPR2a*), or an exon 12-deficient variant (*BMPR2b*) lacking a cytoplasmic tail domain that is unique to BMPR-II among type-II TGFβ superfamily receptors^15,16^. In contrast, the direct interaction of *BMPR2*-derived transcripts with regulators of fundamental cellular processes has received minimal attention^17^. Moreover, the capacity of the *BMPR2* gene to produce alternative transcripts that directly influence cellular function, such as long non-coding or circular RNAs (circRNAs), has not been reported.

CircRNAs are covalently closed, single-stranded RNAs that are resistant to degradation due to the absence of free 5’ or 3’ ends. These transcripts can code for proteins, or regulate the availability, subcellular localization, and functionality of other RNAs or proteins, making circRNAs a promising source for novel regulators of cellular function^18^. Despite this promise, the study of circRNAs is limited by the methods available to both identify and quantify these transcripts in specific cells or tissues. Over 90,000 putative circRNAs have been annotated across the human genome^19^. However, few of these annotations have been validated experimentally and only a fraction of this subset are captured by commercially available circRNA microarrays^20^.

Here, we examine the impact of *BMPR2* loss on the circRNA expression profile of human pulmonary artery endothelial cells (HPAECs). Using ultra-high depth RNA-sequencing and an unbiased screen of 92,369 known or proposed human circRNAs, we identify two novel circRNAs derived from the *BMPR2* locus. In doing so, we define a previously unappreciated interaction between linear and circular *BMPR2* transcripts in the regulation of endothelial proliferation and translational stress responses. Together, these studies not only provide a complete profile of circRNA expression in the human pulmonary endothelium, but also offer an alternative model for endothelial dysfunction in PAH, whereby certain aspects of the diseased cellular phenotype are attributable to the competing actions of linear versus circular *BMPR2* RNA, rather than a loss of the BMPR-II protein product.

## Materials and Methods

### Cell culture

Unless stated otherwise, HPAECs and blood outgrowth endothelial cells (BOECs) were plated at a density of 21,000 cells/cm^2^ and cultured in endothelial growth media (EGM-2) with 2% FBS or EGM-2MV with 10% FBS, respectively (both Lonza, Basel, Switzerland). For inducing quiescence, HPAECs were cultured for 4 hours in endothelial basal medium (EBM-2, Lonza) with 2% FBS. Demographic and mutation data for patient and control BOEC lines can be found in **Table S1.**

### siRNA silencing

Cells were treated with either SMARTpool small interfering RNAs (siRNAs) targeting a range of *BMPR2* transcripts, SMARTpool *CAPRIN1* siRNAs, a non-targeting SMARTpool siRNA control, or custom-designed siRNAs targeting individual circular or linear *BMPR2* transcripts (all Horizon Discovery, Waterbeach, UK). The target sequences for custom siRNAs and catalog information for SMARTpool siRNAs are listed in **Table S2**. For siRNA silencing, cells were cultured in Opti-MEM reduced serum media (ThermoFisher, Waltham, MA) for 3 hours prior to transfection in fresh Opti-MEM containing DharmaFECT 1 transfection reagent (Horizon Discovery) and siRNAs for 4 hours. For studies examining the silencing of a single target (SMARTpool or custom), siRNAs were used at a final concentration of 10 nmol/L. For experiments examining the combined loss of *circ5078* and *CAPRIN1*, 10 nmol/L of each siRNA was used, with non-targeting control siRNA added as needed to bring the final siRNA concentration in all treatment groups to 20 nmol/L.

### RNA isolation and sequencing

One day after plating, HPAECs were transfected with SMARTpool *BMPR2* siRNAs or a non-targeting control. The following day, cells were quiesced for 4 hours, after which they were returned to growth media, with or without 1 ng/mL recombinant human BMP9 (R&D Systems, Minneapolis, MN) for an additional 24 hours. Total RNA was isolated using TRIzol RNA Isolation Reagent (ThermoFisher) and purified using Direct-Zol RNA isolation columns (Zymo Research, Irvine, CA) in accordance with the manufacturer’s instructions, and assessed using a Nanodrop 2000 spectrophotometer (ThermoFisher).

Samples were depleted of ribosomal RNA (NEBNext rRNA Depletion Kit (Human/Mouse/Rat), New England Biolabs, Ipswich, MA) and sequenced using an Illumina NovaSeq 6000 at a mean depth of 312 million paired-end reads per sample (624 million total reads per sample). Reads were trimmed and filtered for quality using Trimmomatic^21^, and the quality of the remaining reads was assessed using FastQC (https://www.bioinformatics.babraham.ac.uk/projects/fastqc/).

CircRNA-derived reads were quantified using methods previously described^22^. Briefly, a publicly available list of known or proposed circRNA annotations within the human genome^19^ was used to construct a library of unique splice junctions for each annotated circRNA. Junctions were constructed by extracting 80 bases to the left and right of each splice site from the linear genome and reassembling these sequences in the proper backspliced orientation. Reads measuring 101 base pairs in length were aligned to the junction library using Bowtie2^23^, with a minimum score setting of -30. Combining all parameters, reads were required to have a minimum overlap of 21 bases on either side of the junction, with a maximum of five mismatched bases across the entire read, in order to be captured. Aligned reads were quantified using featureCounts^24^. Following the filtering of circRNAs for abundance, differential expression analysis was conducted using DESeq2^25^. Publicly available GEO datasets from HPAECs (GSE118446), mouse endothelial cells (GSE216635), and rat whole lung (GSE159668) were profiled for circular and linear *BMPR2*-derived transcripts using a similar approach, with junction size and alignment parameters adjusted in accordance with the sequencing read length. Reverse complementary matches within exons 11 and 12 of the human *BMPR2* gene were identified as described by Ivanov and colleagues^26^.

### qPCR

RNA was reverse transcribed to cDNA using SuperScript IV VILO (ThermoFisher). qPCR analysis was performed using SYBR Green Master mix (Applied Biosystems, Waltham, MA). Relative gene expression was determined using the ΔΔCt method^27^. Primer sequences can be found in **Table S3**. Generic *BMPR2* primers targeted the boundary of exons 6 and 7, which are shared by all linear *BMPR2* transcripts. *BMPR2a* primers targeted the junction between exon 12 (forward) and exon 13 (reverse), while *BMPR2b* primer pairs relied on a forward primer targeting exon 11, and a reverse primer spanning the exon 11/13 splice junction. The amplicon produced by each set of circRNA primers spanned the unique backspliced circRNA junction. As such, circRNA primers align divergently on the linear genome.

### RNase R Digestion

Total RNA (1 μg) from untreated HPAECs was digested by 2U of RNase R enzyme (Lucigen, Middleton, WI) in 10 μL reactions containing 1x digestion buffer. Digestions were performed at 37°C for 20 minutes. 5 μL of digestion product was reverse transcribed and analyzed by qPCR as described above. Samples were compared to a no enzyme control to determine the impact of RNase R digestion.

### Cell proliferation assays

For proliferation assays, HPAECs were plated at a density of 10,500/cm^2^ in 24-well plates and transfected with siRNAs one day after plating. The following day, HPAECs were quiesced for 4 hours before being switched to EGM-2 with 5% FBS at time 0 of the proliferation assay. Cells were trypsinized and counted at 0- and 96-hours, using a hemocytometer and trypan blue staining to determine cell viability. Media was replaced at the 48-hour time point and proliferation was calculated at 96 hours relative to cell number at time 0.

### Protein isolation and immunoblotting

Buffer compositions can be found in **Table S4**. For whole cell lysates, cells were washed with PBS, snap frozen on dry ice, and lysed in RIPA buffer. For cell fractionation lysates, cells were washed in PBS containing 100 μg/mL cycloheximide (CHX, Sigma-Aldrich), scrape collected in homogenization buffer, and passed five times through a 25G needle. Lysates were centrifuged at 14,000g for 10 minutes at 4°C, and the pellet was resuspended in cytoskeletal bound polysome (CBP) buffer, incubated on ice for 30 minutes, and centrifuged again. The pellet was then washed with CBP, resuspended in membrane bound polysome (MBP) buffer, incubated on ice for 10 minutes, centrifuged, and the supernatant removed. Protein concentration was assessed by DC Protein Assay (Bio-Rad, Hercules, CA), as per the manufacturer’s instructions.

Whole cell or cell fraction lysates were run on 10-12% SDS-PAGE gels and transferred to Immobilon-FL PVDF membranes (MilliporeSigma, Burlington, MA). Revert 700 total protein staining (LiCOR, Lincoln, NE) was performed following transfer for normalization within each lane. Following destaining, membranes were blocked in 5% BSA in TBS with 1% Tween (TBST, ThermoFisher) and incubated with primary (overnight) and secondary (1 hour) antibodies in 5% BSA in TBST, as detailed in **Table S5**. Membranes were imaged using the Odyssey Clx Infrared Imaging System and analyzed by Image Studio Software (LiCOR).

### Apoptosis assay

24 hours after siRNA silencing, HPAECs were transferred to quiescence media containing 100 U/mL penicillin, 100 mg/mL streptomycin, and 0.25 mg/mL amphotericin B for 16 hours, followed by an additional 4 hours with or without an apoptotic stimulus of 10 ng/ml TNFα (Peprotech, Cranbury, NJ) and 20 μg/mL CHX. Cells were trypsinized and stained with FITC-conjugated annexin-V and propidium iodide (PI, BD Biosciences, Franklin Lakes, NJ) as per the manufacturer’s instructions. Apoptotic cells were determined by events deemed positive for annexin-V and negative for PI by flow cytometry.

### RNA-Immunoprecipitation

Lysate collection and immunoprecipitation were performed using the Magna RIP immunoprecipitation assay (MilliporeSigma) and 5 μg of rabbit anti-Caprin-1 (Proteintech, Rosemont, IL) or rabbit IgG control, in accordance with the manufacturer’s directions. Protein content of input lysates and immunoprecipitation products were analyzed by immunoblotting as described above. *BMPR2* transcripts from both whole cell lysates and the immunoprecipitated RNA products were amplified by PCR using the primers detailed above and analyzed by agarose gel electrophoresis.

### Sodium arsenite stress assays

Cells were plated on collagen-coated glass chamber slides or 12 mm coverslips and transfected with siRNAs, as described above. One day after siRNA treatment, cells were treated with either fresh EGM-2 or EGM-2 containing 100 μM of sodium arsenite (Sigma-Aldrich) for 1 hour prior to fixation and staining, or lysis and immunoblotting for total and phosphorylated eIF2α, as detailed above. For the assessment of stress granules, cells were washed twice with PBS, fixed with 4% PFA for 15 minutes at room temperature, washed twice in PBS with 0.1% Tween (PBST), blocked in 5% BSA in PBST for 1 hour at room temperature, and labeled overnight at 4°C in 1% BSA in PBST containing mouse anti-G3BP1 (BD Biosciences, 1:1000 dilution) and rabbit anti-Caprin-1 (Proteintech, 1:500). The following day, cells were washed twice with PBST and incubated for 1 hour at room temperature in AF568 donkey anti-mouse and AF647 donkey anti-rabbit (both ThermoFisher, 1:1000) in 1% BSA in PBST. Nuclei were labeled with 600nM DAPI in PBS for 10 minutes at room temperature and slides were mounted in Mowiol. To quantify stress granule positive cells, a minimum of four independent wide-field epifluorescence images were collected per condition, per replicate. For STED imaging of stress granules, cells were labeled with AF532 donkey anti-mouse and AF488 donkey anti-rabbit secondary antibodies (both ThermoFisher, 1:1000) and slides were mounted in ProLong Diamond Antifade with DAPI.

### OPP Assay

Protein synthesis was assessed using the Click-iT Plus OPP protein synthesis kit (ThermoFisher). Briefly, cells were cultured in EGM-2 media, with or without 100 μM of sodium arsenite for 30 minutes prior to incubation with OPP reagent for an additional 30 minutes of culture. Cells were then fixed in 4% PFA and permeabilized with PBST. OPP-containing peptides were labeled with Alexa488, and nuclei labeled with NuclearMask stain as per the manufacturer’s instructions. Fluorescence intensity for both stains was determined using a SpectraMax M3 plate reader (Molecular Devices, San Jose, CA). The nuclear signal was used to normalize the OPP derived signal across wells.

### Polysome Profiling

HPAECs were plated in 15cm dishes and treated with siRNAs, as described above. The day following knockdown, cells were treated with 100 μg/mL of CHX for 15 minutes at 37°C. Following incubation with CHX, cells were scrape collected in ice cold PBS with 100 μg/mL CHX, pelleted at 1,000g for 5 minutes at 4°C, and lysed in 350μL of polysome lysis buffer (**Table S4**). Lysates were incubated on ice for 30 minutes, and insoluble material was pelleted at 13,000g for 10 minutes at 4°C. RNA content was quantified by Nanodrop, and 100 μg of total RNA per condition was loaded onto a 10-45% sucrose gradient (**Table S4**). Gradients were centrifuged at 40,000 rpm for 1.5 hours at 4°C and were fractionated using a BioComp fractionator (BioComp Instruments, Fredericton NB) into 800 μL fractions that were divided equally for the isolation of either protein or RNA. RNA UV absorbance was quantified across the gradient and peak 43/48S, 60S and 80S absorbance values were calculated for each experimental replicate by subtracting background absorbance, determined as the lowest absorbance recorded for the si*Linear+5078* gradient in that replicate.

Protein was isolated by overnight ethanol precipitation at -20°C and centrifugation at 20,000g for 25 minutes at 4°C. Pellets were resuspended in RIPA and 1x loading buffer for immunoblotting. RNA was precipitated from each fraction, as well as from the total polysome lysate input, using sodium acetate and ethanol overnight at -80°C. Precipitates were pelleted at 20,000g for 30 minutes at 4°C and RNA was purified by TRIzol/chloroform extraction for subsequent analysis by qPCR or RNA-sequencing.

For the assessment of translational efficiency, equal volumes of RNA from fractions 8-13 (actively translated RNAs) were pooled for sequencing, alongside samples of total input lysate. Samples were polyA enriched and sequenced, with reads processed as described above. Reads corresponding to transcripts annotated for the GRCh38 genome assembly were quantified using featureCounts. Transcripts not expressed in every sample were filtered out, and changes in RNA expression for the remaining transcripts were determined using DESeq2. Transcripts deemed to be translationally regulated were those exhibiting a differential abundance in polysome fractions relative to the si*Control* treatment (*p_adj_* < 0.05), despite no significant change in abundance in the corresponding total RNA inputs (*p_adj_* ≥ 0.05). Gene set enrichment analysis (GSEA) on selected transcript analysis was performed using Reactome software^28^.

### Seahorse Assay

HPAECs were plated in a 24-well Agilent Seahorse XFe24 cell culture plate at a density of 30,000 cells per well in EGM-2 media and allowed to adhere overnight. The following day, cells were subject to siRNA mediated knockdown as described above. Twenty-four hours after knockdown, oxygen consumption rate (OCR) was determined via the Seahorse XF Mito Stress Test using a Seahorse XFe24 Extracellular Flux Analyzer (Agilent). The sensor cartridge was hydrated in XF Calibrant solution overnight at 37°C. Prior to analysis, cells were washed twice with XF DMEM (supplemented with 1 mM sodium pyruvate, 2 mM L-glutamine, and 25 mM glucose, pH 7.4), and covered in 0.45 mL of fresh XF DMEM for analysis. During the experiment, four baseline measurements were taken, before the sequential addition of 1 μM oligomycin, 1 μM FCCP, and a combined solution containing 1 μM each of rotenone and antimycin A. Three measurements were taken following the addition of each drug or drug cocktail. Following the assay, protein content of each well was determined by a DC Protein Assay (Bio-Rad) for normalization. Following normalization, the following parameters were calculated: non-mitochondrial oxygen consumption (minimum rate following rotenone and antimycin A addition; this was subtracted from all other parameters), basal respiration (final reading prior to oligomycin addition), maximal respiration (maximum reading following FCCP addition) and ATP-coupled respiration (minimum rate following oligomycin, subtracted from basal respiration). These parameters were used to derive spare respiratory capacity (maximal respiration as a percentage of basal respiration) and coupling efficiency (ATP-coupled respiration as a percentage of basal respiration).

### Statistical Analysis

All statistical analyses were performed using Graphpad Prism software. Details of the statistical tests used are specified in the figure captions. Data are presented as mean ± SEM. For work in HPAECs, indicated sample sizes are the number of experimental replicates within the same cell line. For work in BOECs, indicated sample sizes represent the number of individual human subjects tested.

## Results

### Profiling HPAEC circRNA expression identifies BMPR2-derived circRNAs

Global circRNA expression was profiled in *BMPR2* silenced and control HPAECs, with and without treatment with BMP9 as an endothelial-selective BMPR-II ligand^7^. In order to select for the low-abundance reads spanning circRNA backsplice junctions, RNA sequencing was completed at a depth of over 300 million high-quality paired-end reads (600 million individual reads) per sample (**Figure S1**), roughly 10-fold the depth required for standard mRNA differential expression analysis. To profile circRNA expression within this dataset, a junction sequence library was constructed using a publicly available database of 92,369 known or proposed circRNA annotations within the human genome^19^, with junction sequences consisting of the 80 bases to the left and right of the unique backsplice site for each circRNA annotation (**Figure 1A**)^22^. Screening reads against this library filtered out over 99.99% of all reads, leaving only those that aligned with circRNA-derived backsplice junctions (**Figures 1B and 1C**). While 19,351 unique circRNA annotations were detected in at least one sample, an additional filtering of these annotations at a threshold of 0.1 transcripts per kilobase million (TPM) per sample (**Figure 1D**) produced a list of 425 putative circRNAs that are abundantly expressed in HPAECs (**File S1**). These circRNAs represented just over 2% of the annotations, but roughly two thirds of the reads that were captured from the junction library.

**Figure 1.**
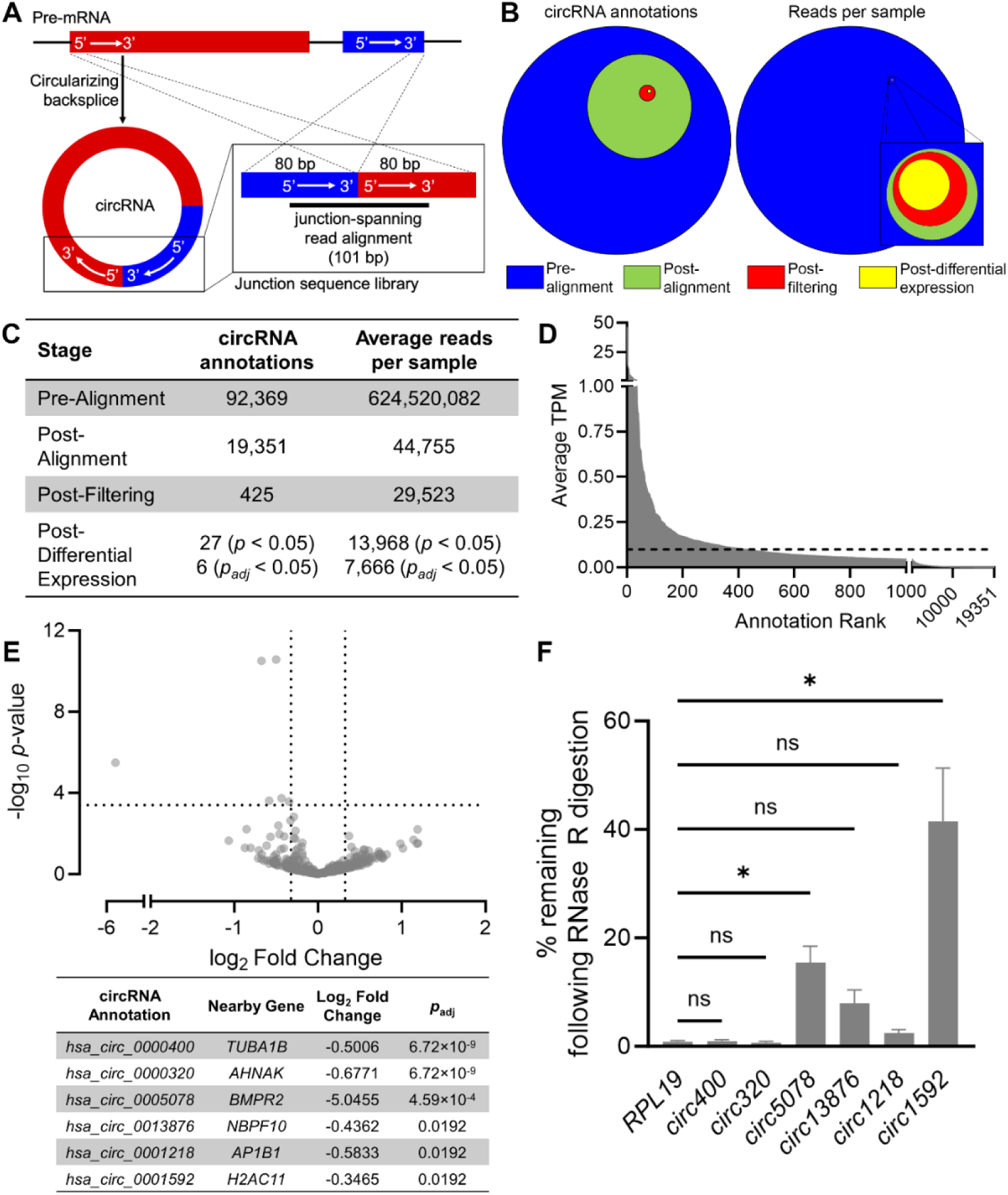
Screening of an ultra-deep RNA-sequencing dataset to profile HPAEC circRNA expression. **(A)** Schematic of bioinformatic analysis method used to identify circRNA-derived sequencing reads. **(B)** Graphical representation and **(C)** summary of the number of circRNA annotations and reads per sample at each stage of analysis. **(D)** circRNA annotations ranked by average transcripts per kilobase million (TPM) across 12 samples. An average TPM ≥ 0.1 was selected as the cut-off of abundantly expressed circRNA annotations, as indicated by the dashed line. **(E)** Volcano plot demonstrating the impact of *BMPR2* silencing on the expression of 425 abundantly expressed circRNAs (red circles in **B**). Annotations with *p_adj_* < 0.05 are summarised. **(F)** RNase R validation of differentially expressed circRNA annotations alongside *RPL19* as linear RNA control (n=7). **(F)** One-way ANOVA with Dunnett’s post-hoc test. ns indicates not significant. \**p* ≤ 0.05.

Analysis of these 425 annotations did not yield any differentially expressed targets with BMP9 treatment. However, 27 differentially expressed annotations were identified with *BMPR2* silencing at a *p*-value of less than 0.05, including six annotations with an adjusted *p*-value less than 0.05 (**Figure 1E and Table S6**). RNase R digestion, which preferentially degrades linear transcripts via their 3’ termini^29^, validated two (*hsa_circ_0005078* and *hsa_circ_0001592*) of these six putative transcripts as true, RNase R resistant circRNAs (**Figure 1F**). Two additional transcripts (*hsa_circ_0013876* and *hsa_circ_0001218*) exhibited partial RNase R resistance relative to the linear RNA control, suggesting the capture of both linear and circular RNA reads for these annotations. Analysis of read alignments for the two annotations that failed RNase R validation (*hsa_circ_0000400* and *hsa_circ_0000320*) identified a bias of these alignments towards segments of their respective junction sequences that were shared with linear mRNA transcripts of the same genomic origin, indicating a misattribution of reads from linear mRNAs (**Figure S2**).

Of the two validated circRNA annotations from the differential expression analysis, *hsa_circ_0005078* (*circ5078*) was of particular interest, as it consists of a circularized form of exon 12 from the *BMPR2* locus (**Figure 2A**). A subsequent examination of the 19,351 annotations arising from the junction library screen identified an additional seven *BMPR2*-derived annotations that were detected in at least one sample, of which one, *hsa_circ_0003218* (*circ3218*), was also abundantly expressed in HPAECs (**Figure 2B**). Like *circ5078*, *circ3218* was resistant to RNase R digestion (**Figure 2C**). However, while *circ5078* was targeted by the *BMPR2* siRNA pool used in the RNA-sequencing screen (**Figures 2A and 2D**), *circ3218*, which consists of *BMPR2* exons 2 and 3, was unaffected by *BMPR2* siRNA treatment (**Figures 2A and 2E**). An independent validation using lower-depth, publicly available GEO datasets confirmed the existence of *BMPR2b*, *circ5078* and *circ3218* in human, but not mouse endothelium (**Figure S3**). Similarly, *circ3218* was the only alternative *BMPR2* transcript to be detected in the rat lung (**Figure 2F**), suggesting that the splicing events responsible for *BMPR2b* and *circ5078* formation do not occur in rodents. This finding was supported by an analysis of the intronic sequences flanking exon 12 in the human *BMPR2* gene. A screen for the reverse complementary matches (RCMs) that promote pre-mRNA hairpin formation and circularization identified multiple RCMs between a single region of intron 11 and seven distinct sites within intron 12 (**Figure 2G**). All but one of these RCMs were identified as ALU family repetitive elements, which are known to be enriched in the introns flanking human circRNAs (**Figure 2H**)^30^. Importantly, while some of these RCMs were conserved in Rhesus macaques, none were conserved in mice or rats, explaining the absence of *circ5078* biogenesis in those species.

**Figure 2.**
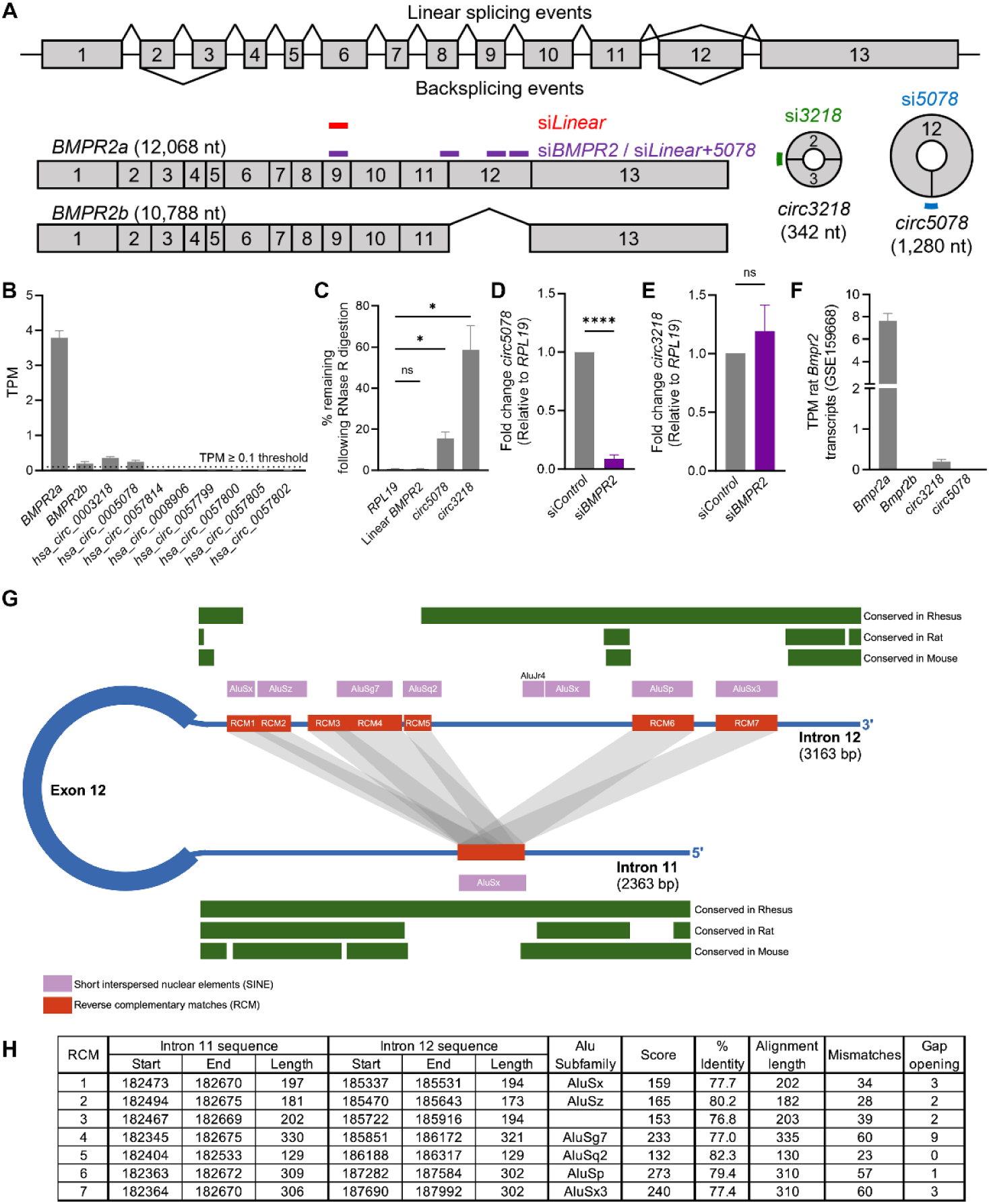
Identification of novel *BMPR2* derived circRNAs. **(A)** Summary of all linear and circular *BMPR2*-derived transcripts expressed in HPAECs. Regions targeted by siRNAs used in this study are indicated. **(B)** Relative expression of linear *BMPR2* transcripts and the 8 *BMPR2*-derived circRNA annotations that were detected from the RNA-seq screen (n=12). Dashed line indicates expression cutoff of 0.1 TPM. **(C)** RNase R validation of *BMPR2*-derived circRNA annotations, alongside linear *BMPR2* mRNA and *RPL19* as a linear RNA control (n=6). **(D)** Relative expression of *circ5078* and **(E)** *circ3218* in HPAECs treated with pooled siRNAs targeting *BMPR2* (si*BMPR2*) or a non-targeting siRNA control pool (si*Control*) (n=5). **(F)** Quantification of human *BMPR2* transcript equivalents in RNA-sequencing data from rat lungs (GSE159668, n=6). **(G)** Graphical depiction of the introns surrounding exon 12 of *BMPR2*, the reverse complementary matches (RCMs) they contain, and their conservation in rodent and non-human primate species. **(H)** Summary of the alignments between the RCM sequences within intron 12 of *BMPPR2* and the indicated segment of intron 11. **(C)** One-way ANOVA with Dunnett’s post hoc test. **(D, E)** Paired t-test. ns indicates not significant. \**p* ≤ 0.05, \*\*\*\**p* < 0.0001.

### BMPR2 circRNAs regulate endothelial apoptosis and proliferation

To investigate the functional contribution of these novel *BMPR2* transcripts to endothelial function, transcript-specific siRNAs were designed to selectively deplete linear *BMPR2* mRNAs (si*Linear*), *circ5078* (si*5078*), or *circ3218* (si*3218*), alongside the commercially available *BMPR2* siRNA pool (si*Linear*+*5078*) that targets both linear *BMPR2* transcripts and *circ5078* (**Figure 3A**). Although both circRNA-specific siRNAs modestly reduced linear *BMPR2* mRNA levels, neither siRNA impacted BMPR-II protein, which was reduced exclusively in the si*Linear* and si*Linear+5078* groups (**Figures 3B and 3C**). Functionally, the silencing of *circ3218*, but not *circ5078*, increased the susceptibility of HPAECs to TNFα and cycloheximide-induced apoptosis (**Figures 3D and 3E**). In contrast, *circ5078* depletion significantly enhanced HPAEC proliferation when compared to the non-targeting siRNA control (**Figure 3F**). Interestingly, the combined silencing of *circ5078* and linear *BMPR2* mRNAs eliminated the proliferative effect achieved by silencing *circ5078* alone, suggesting a potential interaction between *circ5078* and linear *BMPR2* mRNA in the regulation of HPAEC proliferation.

**Figure 3.**
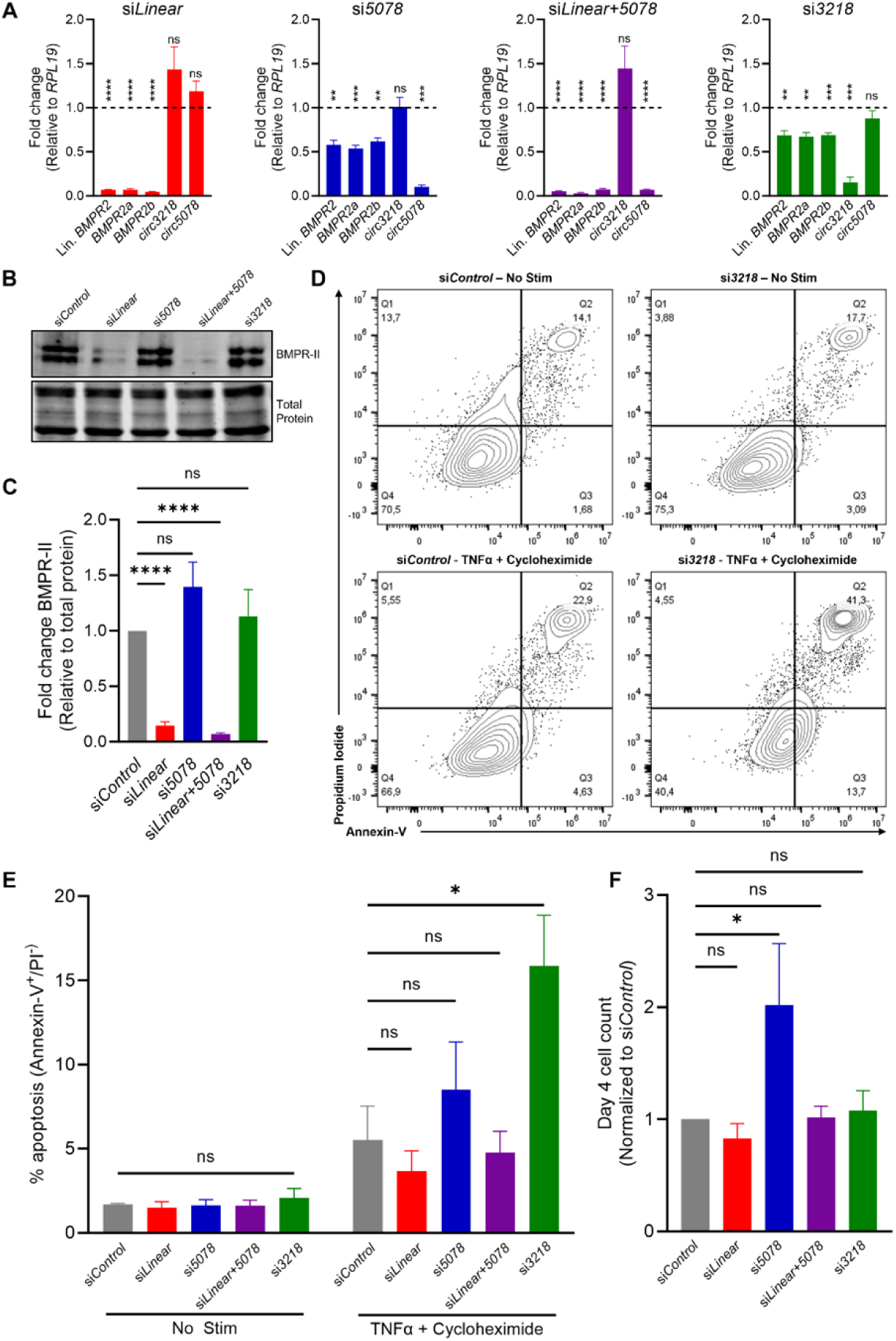
Functional characterization of *BMPR2*-derived circRNAs. **(A)** Impact of si*Linear* (n=4), si*5078* (n=6), si*Linear+5078* (n=5) and si*3218* (n=6) on the abundance of generic linear *BMPR2*, and individual *BMPR2* transcripts. Dashed line represents the expression of each transcript in the si*Control* sample, bars represent the expression of each transcript under the respective knockdown condition. **(B)** Representative immunoblot and **(C)** quantification of BMPR-II protein expression under each knockdown condition (n=7). **(D)** Representative flow plots of propidium iodide (PI) and FITC-annexin-V stained HPAECs and **(E)** quantification of Annexin-V^+^/PI^-^ apoptotic cells under each siRNA condition, with and without a TNFα and cycloheximide apoptotic stimulus (n=7). **(F)** Relative change in cell number over 96 hours for each siRNA treatment relative to si*Control* from experiments in **A**. **(A)** Two-way ANOVA with Bonferroni post hoc test. **(C)** One-way ANOVA with Dunnett’s post hoc test. **(E)** Two-way ANOVA with Dunnett’s post hoc test. **(F)** Mixed effects model with Dunnett’s post hoc test. ns indicates not significant. \**p* ≤ 0.05, \*\**p* < 0.01, \*\*\**p* < 0.001, \*\*\*\**p* < 0.0001.

### The proliferative effect of circ5078 loss is Caprin-1-dependent

An *in silico* BEAM RNA Interaction mOtifs (BRIO)^31^ screen for potential *circ5078* protein targets identified three likely interacting partners for *circ5078*, including Argonaute 2, Nuclear Cap Binding Subunit 3 (NCBP3) and Caprin-1 (**Figure 4A**). Of these targets, Caprin-1 was of particular interest as a major component of cytoplasmic RNA granules that is known to enhance proliferation via the increased translation of cell cycle regulators^32^. RNA-immunoprecipitation for Caprin-1 enriched both *circ5078* and linear *BMPR2* mRNA in HPAEC lysates (**Figure 4B**). Although *circ5078* was not amenable to visualization by custom fluorescence *in situ* hybridization (FISH) using RNA probes targeting its unique backsplice junction, linear *BMPR2* mRNA targeting FISH probes were found to co-localize with Caprin-1 by super-resolution microscopy, both at baseline and within arsenite-induced stress granules (**Figure S4**).

**Figure 4.**
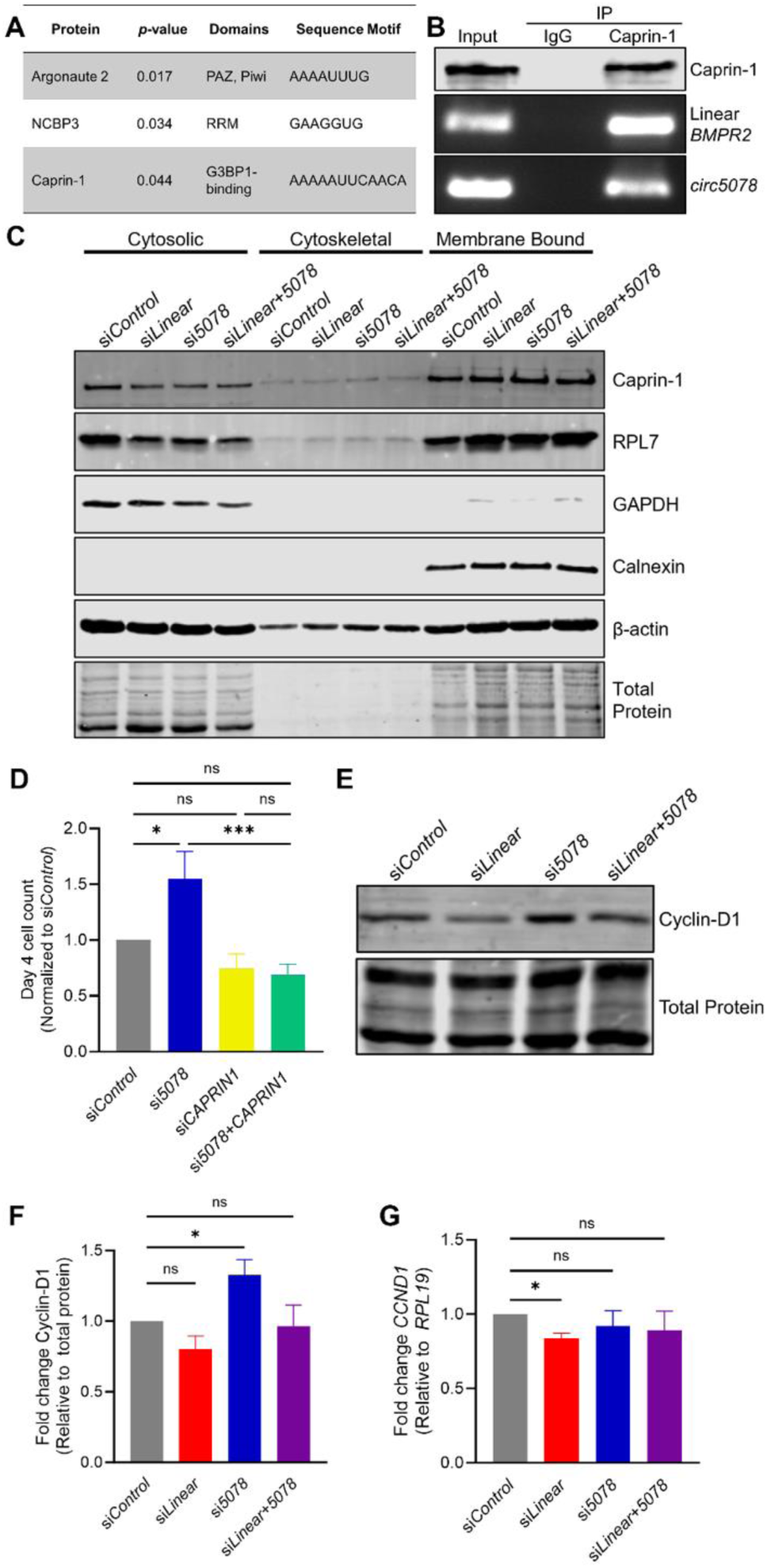
Regulation of proliferation by *circ5078* is Caprin-1 dependent. **(A)** Potential *circ5078* protein binding partners identified by BRIO. **(B)** RNA-immunoprecipitation of linear *BMPR2* mRNA and *circ5078* with anti-Caprin-1 antibody or IgG control. **(C)** Representative immunoblot demonstrating Caprin-1 protein abundance in cytosolic, cytoskeletal, and membrane bound fractions following treatment with siRNAs targeting linear *BMPR2* mRNA, *circ5078*, or both, versus non-targeting si*Control*. **(D)** Relative change in cell number over 96 hours of HPAECs following silencing of *circ5078*, *CAPRIN1*, or both RNAs, relative to si*Control* (n=8). **(E)** Representative immunoblot and **(F)** quantification of Cyclin-D1 protein following treatment with si*Control*, si*Linear*, si*5078*, or si*Linear+5078* siRNAs (n=11). **(G)** *CCND1* gene expression under each siRNA condition (n=5). **(D)** One-way ANOVA with Bonferroni post hoc test. **(F, G)** One-way ANOVA with Dunnett’s post hoc test. ns indicates not significant. \**p* ≤ 0.05, \*\*\**p* < 0.001.

Depletion of *circ5078* or linear *BMPR2* mRNA had no impact on the abundance of Caprin-1 protein or its distribution between the cytosol and a membrane-bound fraction containing the endoplasmic reticulum, the two major sites of protein synthesis in the cell (**Figure 4C**). However, *CAPRIN1* silencing did eliminate the excessive proliferation observed in *circ5078*-depleted HPAECs (**Figures 4D and S5**), confirming that this proliferative response is indeed Caprin-1 dependent. Moreover, *circ5078* depletion induced a significant increase in protein levels of the known Caprin-1 translational target, Cyclin-D1 (**Figures 4E and 4F**), despite no elevation in corresponding mRNA levels (**Figure 4G**). As with the proliferative response, this elevation in Cyclin-D1 protein was lost when *circ5078* and linear *BMPR2* mRNA were silenced together, further supporting opposing roles for linear *BMPR2* mRNA and *circ5078* in the regulation of endothelial proliferation and translational control.

### BMPR2 transcripts differentially regulate stress granule assembly and structure

In keeping with previous reports demonstrating a relationship between *BMPR2* loss and impaired endothelial stress responses^10^, linear *BMPR2* mRNA silencing reduced arsenite-induced stress granule formation, as determined by Caprin-1 and G3BP-1 co-localization in HPAECs (**Figures 5A-C**). *circ5078* depletion had the opposite effect, enhancing both the percentage of granule-positive cells, as well as the size, density, and abundance of stress granules in these cells (**Figures 5B-C and S6**). As with the proliferation studies, pooled silencing of linear *BMPR2* mRNA and *circ5078* restored arsenite-induced stress granule formation to control levels, despite the complete absence of BMPR-II protein in this experimental condition. Although the impairment of stress responses with *BMPR2* loss was previously linked to reduced eIF2α phosphorylation^10^, phosphorylated eIF2α was unchanged with *BMPR2* transcript depletion at baseline and was induced to an equivalent degree in all siRNA treatment groups in response to arsenite-induced oxidative stress (**Figures 5D and 5E**).

**Figure 5.**
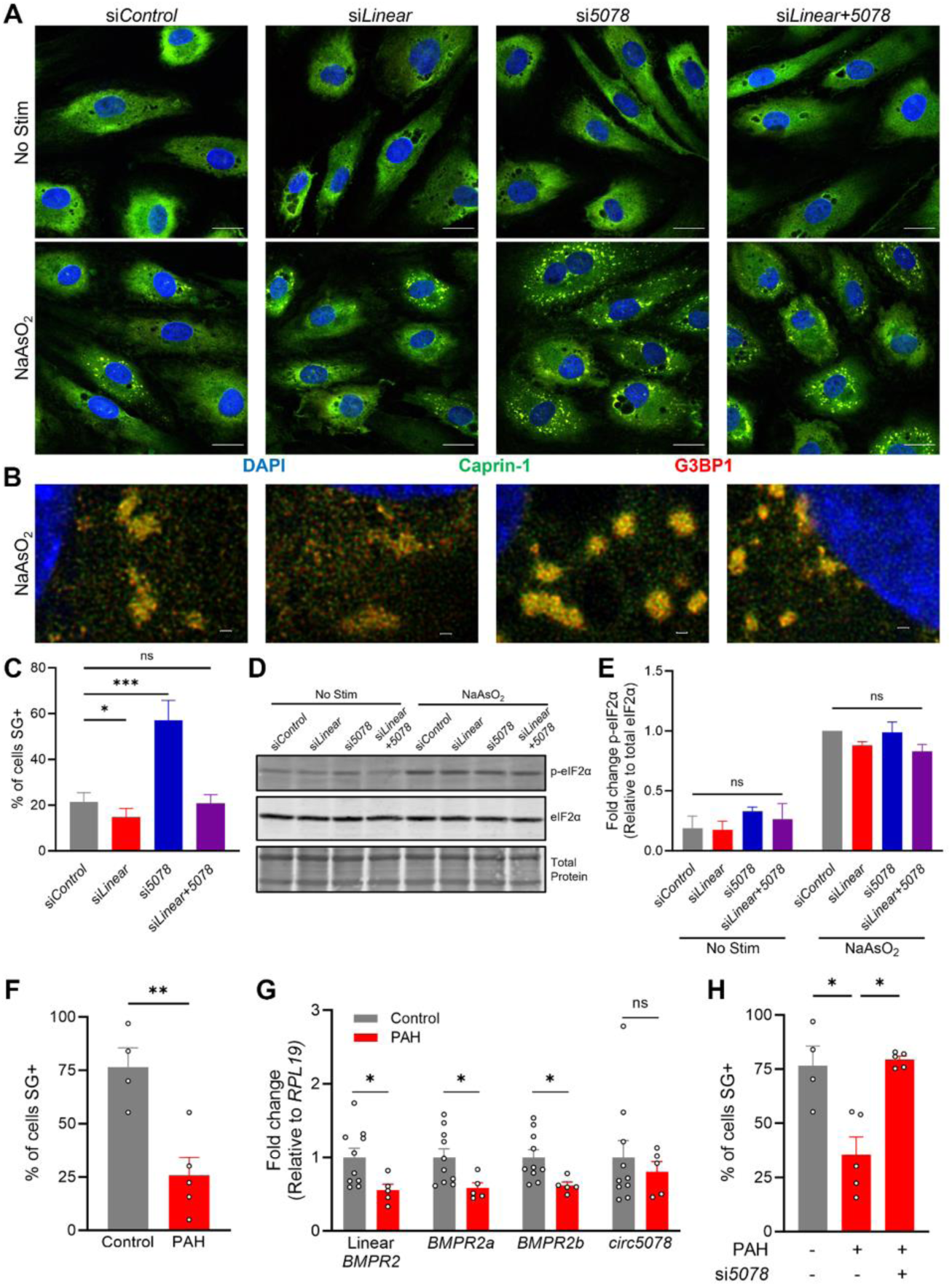
Linear and circular *BMPR2* transcripts differentially modulate Caprin-1 granule formation. **(A)** Representative confocal (Scale bar = 20 μm), and **(B)** STED super-resolution (Scale bar = 0.5 μm) images of HPAECs following 1 hour of treatment with 100 μM sodium arsenite-induced, stained with Caprin-1 (green) and the stress granule marker G3BP-1 (red) under following si*Control*, si*Linear*, si*5078*, or si*Linear+5078* treatment. **(C)** Quantification of stress granules under each knockdown condition (n=9). **(D)** Representative immunoblot and **(E)** quantification of phosphorylated eIF2α in HPAECs (n=3). **(F)** Quantification of cells with stress granules in blood outgrowth endothelial cells (BOECs) from healthy controls (n=4) and PAH patients with *BMPR2* mutations (n=5) in response to sodium arsenite. **(G)** Relative expression of linear *BMPR2*, *BMPR2a*, *BMPR2b*, and *circ5078* transcripts in blood outgrowth endothelial cells from *BMPR2* mutation-bearing PAH patients (n=5) and healthy controls (n=10). **(H)** Stress granule production in control BOECs treated with si*Control* (n=4), and PAH BOECs treated with si*Control* or si*5078* (n=5). **(C, E and H)** One-way ANOVA with Dunnett’s post hoc test. **(F, G)** Unpaired t-test. ns indicates not significant. \**p* ≤ 0.05, \*\**p* < 0.01, \*\*\**p* < 0.001.

Stress granule formation was also impaired in blood outgrowth endothelial cells (BOECs) from *BMPR2* mutation-bearing PAH patients relative to healthy controls (**Figures 5F and S7A**), which was in keeping with a reduction of linear *BMPR2* transcripts, but not *circ5078*, in these cells (**Figure 5G**). More critically, *circ5078* depletion rescued this phenotype in PAH patient BOECs (**Figures 5H and S7B**), indicating that rebalancing the relative abundance of circular to linear *BMPR2* transcripts can restore healthy stress responses in the diseased endothelium, independent of BMPR-II protein levels.

### BMPR2 transcript depletion does not influence global protein synthesis

To determine the impact of *circ5078* and linear *BMPR2* depletion on endothelial translation, lysates from each silencing condition were assessed by sucrose gradient ultracentrifugation (**Figure 6A**), which allowed for the quantification of ribosomal subunits (Fractions 2-4), 80S monosomes (Fraction 5), and actively translating polysomes (Fractions 6-13), as well as the co-sedimentation of the proteins (**Figure 6B**) and RNAs associated with these complexes. Gradient analysis identified an accumulation of free ribosomal subunits and polysome half-mers in control HPAECs (**Figures 6C-E**). This accumulation was not linked to impaired ribosome biogenesis (**Figure S8**), but was eliminated by the silencing of linear *BMPR2* mRNA, *circ5078*, or the pooled depletion of both transcripts. A similar pattern was observed for Caprin-1, which was elevated in fraction 2 in control HPAECs, but reduced with the depletion of *BMPR2*-derived transcripts (**Figure 6B**).

**Figure 6.**
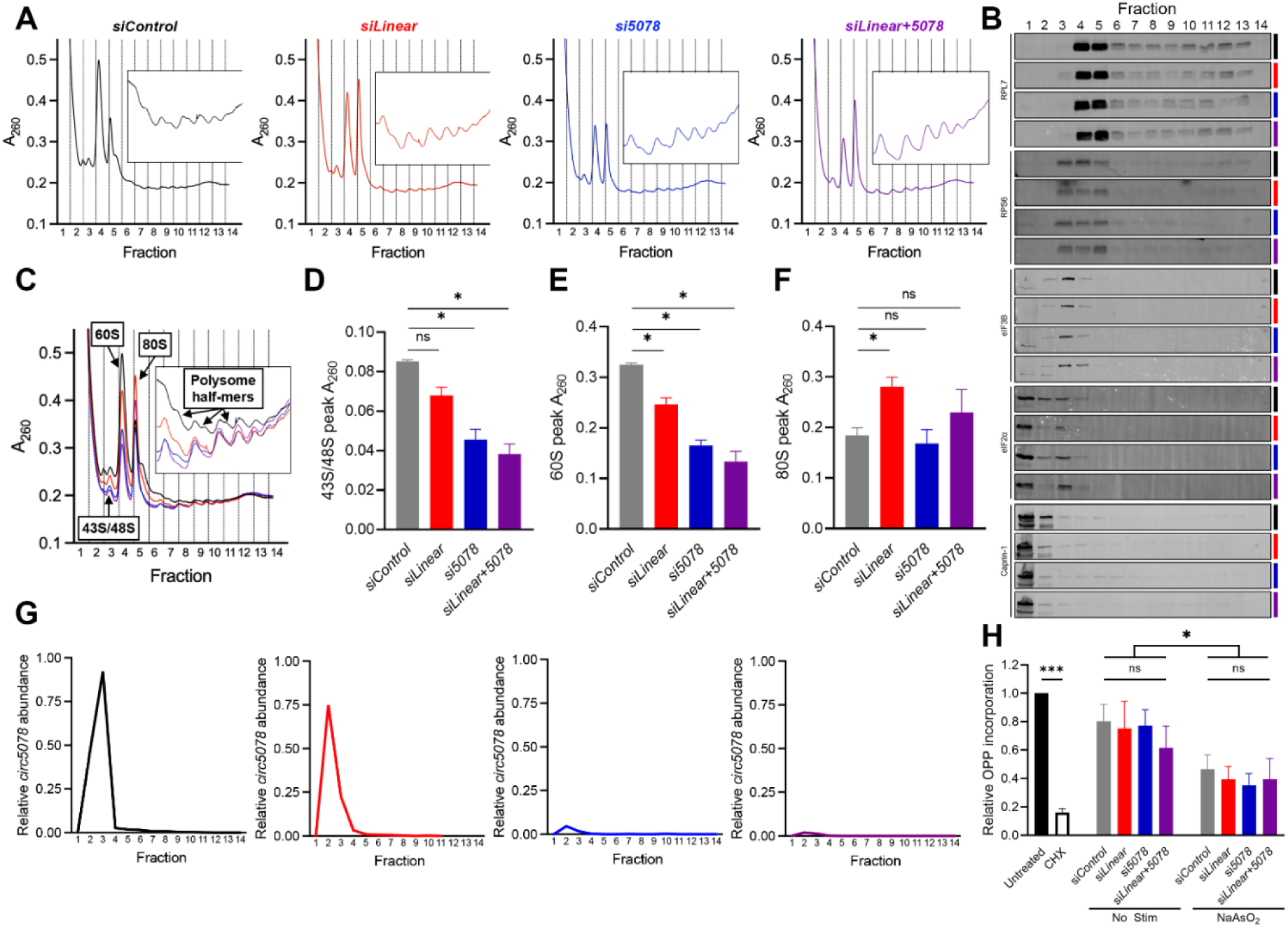
*BMPR2* transcripts regulate ribosomal assembly, but not global protein synthesis. **(A)** Sucrose gradient UV absorbance (A_260_) profiles for HPAECs treated with si*Control* (black), si*Linear* (red), si*5078* (blue), or si*Linear+5078* (purple) siRNAs. Insets are magnification of the curves in fractions 6-12. **(B)** Representative immunoblots of RPL7, RPS6, eIF3B, eIF2α and Caprin-1 across gradient fractions. **(C)** Overlaid A_260_ profiles highlighting the 43/48S preinitiation complex (PIC), 60S ribosomal subunits, 80S monosomes and polysome half-mers. **(D)** Quantification of absorbance less background for the 43S/48S, **(E)** 60S, and **(F)** 80S peaks (n=3 each). **(G)** Relative abundance of *circ5078* RNA across all fractions in each siRNA condition. Data shown is the pooled average of 3 independent experiments. **(H)** Quantification of global protein production by incorporation of O-propargyl-puromycin (OPP) into nascent peptides in HPAECs (n=6). **(D-F)** One-way ANOVA with Dunnett’s post hoc test. **(H)** Paired t-test between untreated and cycloheximide groups. Two-way ANOVA with Dunnett’s post hoc test between all siRNA treated groups. ns indicates not significant. \**p* ≤ 0.05, **\*\*\****p* < 0.001.

In contrast, assessment of the 80S peak identified an increase in monosomes exclusively with linear *BMPR2* mRNA silencing, either alone or in the pooled condition (**Figure 6F**). This increase was accompanied by a potential loss of free eIF2α in fraction 2, versus preinitiation complex (PIC)-associated eIF2α in fraction 3 (**Figure 6B**). While the co-sedimentation of *circ5078* with these fractions further supported a possible role for *BMPR2*-derived transcripts in translational initiation (**Figure 6G**), total protein synthesis was not altered under any of the siRNA conditions, either at baseline or in response to sodium arsenite (**Figures 6H and S9**). Moreover, a higher resolution analysis of initiation complexes, involving the cross linking of samples prior to sedimentation on shallow sucrose gradients, identified no differences in eIF2α distribution with *BMPR2* transcript depletion (**Figures S10A and S10B**). *circ5078*, which was elevated in fraction 7 of these gradients with linear *BMPR2* mRNA depletion (**Figure S10C**), did not co-sediment with components of the PIC or Caprin-1, and no differences were observed in the abundance of free ribosomal subunits or Caprin-1 protein with *circ5078* or linear *BMPR2* mRNA silencing (**Figure S10D**). Together, these discrepancies between the steep polysome gradients and the cross-linked shallow gradients suggest that *BMPR2* transcripts are not regulating global translational initiation, but may instead influence the assembly and stability of ribosomes or translational complexes involved in the translation of specific mRNAs.

### circ5078 regulates the translation of mitochondrial ribosomes and mitochondrial function

To identify if specific mRNAs are differentially translated with *BMPR2* transcript depletion, we sequenced polyadenylated RNA from actively translated fractions of the polysome gradients (Fractions 8-13) for each siRNA condition. Total polyadenylated RNA from input lysates was also sequenced to control for changes in overall transcript abundance arising from siRNA treatment (**Figure 7A**). For each siRNA condition, differential expression analysis was performed on both the polysome and total input samples, relative to their respective non-targeting siRNA controls (**Figure S11**). Differentially translated genes were identified as those not altered in the input samples (*p_adj_* ≥ 0.05), but exhibiting differential abundance in the polysome fraction (*p_adj_* < 0.05) (**Figure 7B and File S2**). This analysis identified 1,137 differentially translated mRNAs across all treatment groups.

**Figure 7.**
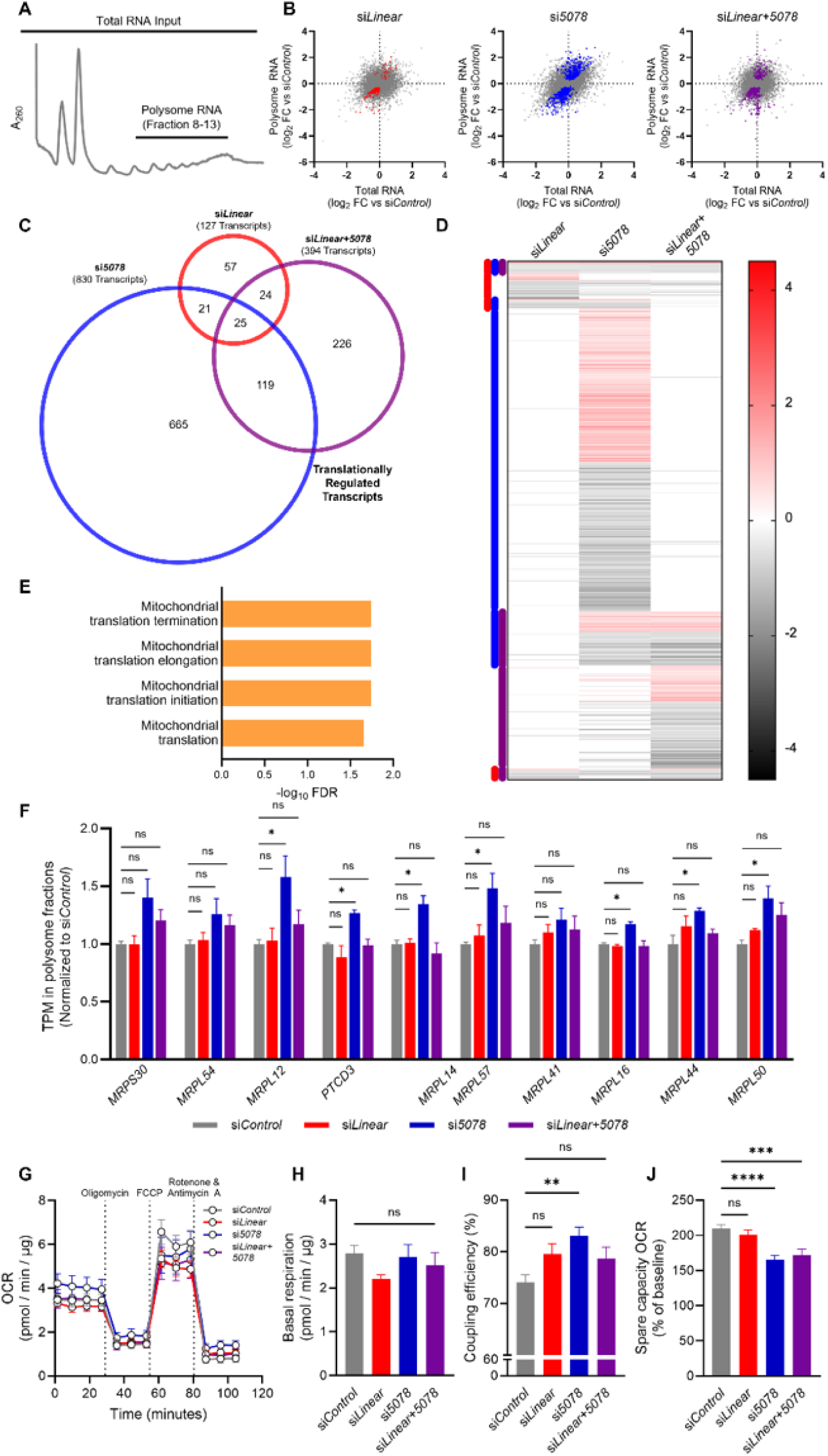
Loss of *circ5078* in HPAECs alters mitochondrial function via altered translation of mitochondrial ribosomes. **(A)** RNA sequencing was performed on total input RNA from unfractionated lysates, and RNA from the actively translated Fractions (8-13) of polysome profiles for HPAECs treated with si*Control*, si*Linear*, si*5078*, or si*Linear+5078* siRNAs. n=3 independent experiments. **(B)** Scatter plots of the log_2_ fold change versus si*Control* in both total RNA and polysome RNA fractions for each siRNA condition. Highlighted transcripts are those deemed to be translationally regulated (*p_adj_* ≥ 0.05 in input fraction, *p_adj_* < 0.05 in polysome fraction). **(C)** Venn diagram and **(D)** heatmap of genes regulated by translation in at least one of the three conditions. **(E)** Summary of pathways enriched among the transcripts in which translation was upregulated by *circ5078* loss alone, as identified by Reactome. **(F)** Relative expression in polysome fractions of transcripts encoding components of the mitochondrial translation pathway. **(G)** HPAECs treated with si*Control* (n=14), si*Linear* (n=12), si*5078* (n=12) or si*Linear+5078* (n=11) were analyzed by Seahorse XFe24 Extracellular Flux Analyzer. Oxygen consumption rate (OCR) was assessed across the course of the experiment following the addition of oligomycin, FCCP, and rotenone and antimycin A. **(H)** Basal respiration, **(I)** coupling efficiency, and **(J)** spare capacity following these treatments. **(F, H-J)** One-way ANOVA with Dunnett’s post hoc test. ns indicates not significant. \**p* ≤ 0.05, \*\**p* < 0.01, \*\*\**p* < 0.001, \*\*\*\**p* < 0.0001.

Of these transcripts, few were shared between the individual siRNA treatments, and more than half were unique to *circ5078* silencing (**Figures 7C**). Although the 665 transcripts impacted by *circ5078* were split evenly between those that were enriched in polysomes, and those that were under-represented in these fractions (**Figure 7D**), gene set enrichment analysis (GSEA) using Reactome^28^ did not identify any pathways that were translationally repressed by *circ5078* depletion. Instead, this analysis identified an upregulation of pathways regulating mitochondrial translational processes (**Figure 7E**), driven by the enrichment of 10 nuclear genes encoding mitochondrial ribosome subunits. A subsequent comparison of transcript abundance in the polysome fraction across all four siRNA conditions confirmed that 7 of these 10 targets were uniquely upregulated with *circ5078* depletion (**Figure 7F**).

Assessment of mitochondrial oxidative metabolism (**Figure 7G**) did not show any significant differences in baseline mitochondrial oxygen consumption rate (OCR) with *BMPR2* transcript silencing (**Figure 7H**). However, coupling efficiency, a measure of mitochondrial ATP production per unit of oxygen consumed, was elevated with *circ5078* depletion (**Figure 7I**). Spare capacity, defined as a cell’s ability to respond to metabolic stress, and calculated as the percentage difference between maximal and baseline OCR, was also reduced with *circ5078* silencing, either alone or in the pooled siRNA condition (**Figure 7J**), providing a potential link between mitochondrial function and the hyperproliferative, stress-sensitive endothelial phenotype of *circ5078* loss.

## Discussion

By identifying *circ5078* as a novel *BMPR2*-derived functional RNA and defining its role as a regulator of endothelial proliferation and stress responses, this study offers the prospect that certain elements of the endothelial PAH phenotype, which were previously attributed to a loss of the BMPR-II protein receptor^10^, may instead arise through an imbalance in the functional effects of distinct *BMPR2* gene products. The capacity of *circ5078* depletion to normalize stress granule formation in PAH patient BOECs, without targeting BMP signaling or restoring BMPR-II protein levels, indicates that loss of the protein receptor may be necessary, but is not sufficient to drive this aspect of the diseased endothelium. Importantly, it is not yet known whether the opposing actions of linear *BMPR2* mRNA and *circ5078* are mediated by competition of the two transcripts for shared binding partners, or whether the effects of linear *BMPR2* mRNA occur through a distinct mechanism that is dependent on the BMPR-II protein product or its receptor kinase activity. Additional studies, involving the rescue of patient BOECs with linear *BMPR2* transcripts encoding either wildtype, exon 12-deficient, or kinase-dead BMPR-II protein would help to resolve this question.

Our work identified Caprin-1 as a potential binding partner for *circ5078* and demonstrated a critical role for this protein in the proliferative phenotype of *circ5078*-deficient HPAECs. As a known regulator of cell cycle^33^, and a central component of both stress granules and the RNA-rich, phase-separated cytoplasmic aggregates that facilitate mRNA translation^34^, Caprin-1 represents an appealing target for understanding how *BMPR2* transcripts influence endothelial function. Although both linear *BMPR2* RNA and *circ5078* were enriched by RNA-immunoprecipitation of Caprin-1, this protein did not co-sediment with *circ5078* when fixed cell lysates were applied to shallow sucrose gradients. This finding suggests that the interaction of *BMPR2* transcripts with Caprin-1 may not involve direct binding, but may instead consist of an indirect association involving other components of these phase-separated Caprin-1-rich granules^34^, such as the RNA-binding protein FMRP^35^, which has been shown to bind *BMPR2* transcripts via multiple sites within exon 12^17^. Additional studies, such as PAR-CLIP immunoprecipitation^36,37^ or an analysis of proteins co-precipitating with a pull-down of circular or linear *BMPR2* transcripts, are needed to determine whether direct *BMPR2* transcript/Caprin-1 interactions exist and to identify other potential *circ5078* interacting partners.

The observation that silencing *BMPR2*-derived transcripts modulates stress granule formation without altering eIF2α phosphorylation differs from previous work reporting a reduction in stress-induced eIF2α phosphorylation in *BMPR2*-deficient endothelial cells via increased GADD34–PP1 phosphatase activity^10^. It is important to note that the absence of differential eIF2α phosphorylation in our studies may be dependent on the nature of the cellular stress employed, as the previous work focused primarily on TNFα-induced stress responses^10^. Caprin-1 deletion and overexpression studies have also shown that, despite the fact that stress granule formation and eIF2α phosphorylation are both central features of the integrated stress response, granule assembly can occur in a manner that is independent of eIF2α phosphorylation or altered translational initiation^32^. When coupled with the observation *BMPR2* transcript depletion did not alter global protein synthesis or the overall assembly of translational initiation complexes, the current work indicates that the loss of *circ5078* does not induce a conventional stress response, but may instead influence endothelial function through the differential translation of specific mRNA transcripts.

Assessment of translationally enriched mRNA transcripts identified a number of mitoribosomal genes that are enhanced exclusively with *circ5078* depletion. Imbalance in mitochondrial versus nuclear-encoded mitochondrial proteins, known as the mitonuclear imbalance, has been reported in both disease and ageing^38^. The identification of increased mitochondrial coupling efficiency specifically in *circ5078*-deficient HPAECs indicates a greater capacity of these cells to use oxygen to generate ATP, providing a potential mechanistic link between endothelial metabolic function and excessive proliferation. The reduction in mitochondrial spare capacity with *circ5078* silencing also supports a mechanism by which *circ5078*-deficient HPAECs are more susceptible to stress. However, the precise integration of these effects with the opposing actions of linear *BMPR2* mRNA has yet to be determined.

This study used high depth RNA sequencing to identify and characterize two novel circRNAs derived from the *BMPR2* gene locus. The approach of screening such a dataset for backsplice junction-spanning reads offers several advantages over alternative circRNA profiling methods, such as RNase R pre-treatment of isolated RNA, or the probing of total RNA isolates using commercially available circRNA microarrays^39,40^. While the infrequency of junction-spanning reads within conventional RNA-sequencing datasets requires extremely high read depths to achieve the statistical power needed for differential expression analysis, the absence of enzymatic digestion steps in this protocol minimizes sample degradation and offers the advantage of evaluating circRNAs alongside traditional linear transcripts. Unlike microarrays, the quantification of circRNA annotations using a backsplice junction library also eliminates the limitation of examining a predetermined set of known transcripts, allowing for the identification of new circRNAs from unvalidated annotations and the retrospective analysis of existing datasets.

In addition to the identification and functional characterization of *circ5078* and *circ3218*, our unbiased junction library screen produced a complete profile of the 425 circRNA annotations that are abundantly expressed in the human pulmonary endothelium. While this profile can serve as a useful tool for understanding the contribution of circRNAs to endothelial cell function in both health and disease, it is critical that any proposed circRNA annotations be validated as true circular transcripts prior to further functional characterization. The importance of proper RNase R validation is highlighted by the fact that two of the six differentially expressed annotations from the original junction library screen were found to arise through the misattribution of reads from linear mRNAs with shared sequences. One of these targets, a proposed circRNA from a portion of exon 5 of the *AHNAK* gene (*hsa_circ_0000320*), was reported as a regulator of triple-negative breast cancer^41^. Further study is now required to determine if this annotation does indeed represent a true circRNA or if its proposed effects are instead attributable to the protein product of its parent gene.

An additional limitation of the current study is the fact that the human-specific nature of *circ5078* precludes more detailed assessments into its role in regulating pulmonary vascular function in rodents, including standard pre-clinical models of PAH. The absence of *circ5078* and *BMPR2b* transcripts in mouse and rat-derived datasets, which we attribute to the absence of reverse complementary sequences within the introns flanking exon 12 in these species, may help to explain why many rodent models of *BMPR2* loss fail to recapitulate the human PAH phenotype or develop a more severe phenotype in response to disease-inducing stimuli. The absence of both *circ5078* and the truncated *BMPR2b* transcriptional variant in rodents also suggests that these transcripts are generated through a shared process involving the excision of exon 12 from full-length *BMPR2* pre-mRNA. While *BMPR2b* expression is elevated relative to that of *BMPR2a* in PAH patients with *BMPR2* mutations when compared to unaffected mutation carriers^42^, *circ5078* expression in this cohort is unknown and merits further investigation.

Overall, our study identifies two new functional *BMPR2* gene products, and in doing so, provides important new insights into the potential of the *BMPR2* genomic locus to regulate the pulmonary endothelium. While we have established novel contributions of both *circ5078* and linear *BMPR2* mRNA to endothelial proliferation, translation, and stress responses, this work also raises many new questions that each require a detailed follow-up investigation. Future work exploring the precise mechanisms by which *circ5078* contributes to endothelial translational control will help to clarify any potential convergence between *EIF2AK4* and *BMPR2* mutations in the regulation of endothelial function, as well as the role of the integrated stress response in pulmonary vascular disease.

## Data Availability

Data supporting this work is available upon request from the corresponding author, MLO. Sequencing data will be made publicly available upon the publication of this manuscript.

## Funding

This work was supported by funding from the Canadian Institutes of Health Research (CIHR; PJT-178078). MMV would like to acknowledge graduate scholarship support from CIHR and the Canadian Vascular Network.

## Acknowledgements

The authors would like to acknowledge Ariana Bacelar-Jimenez, Dr. Lindsey Hawke and Dr. Kathrin Tyryshkin for their technical support, as well as Dr. Lynne Postovit for assistance with the sucrose gradient fractionation studies.

## Notes

### Competing Interest Statement

The authors have declared no competing interest.

### Summary of Updates

This version of the manuscript includes substantial new data, as summarized below: i.A sequence analysis of the human BMPR2 gene, identifying reverse complementary ALU elements in the introns flanking the sequence that gives rise to circ5078. These elements are known to promote RNA circularization, but are not conserved in the rat or mouse genomes ii.A higher resolution assessment of translational initiation complexes in primary pulmonary endothelial cells, with and without BMPR2 transcript depletion, using crosslinked samples separated on shallow (7.5-30%) sucrose gradients iii.Quantification of 5.8S, 18S, and 28S ribosomal RNAs, as well as RPL7 and RPS6 protein expression, addressing the potential for impaired ribosome biogenesis with circ5078 or linear BMPR2 transcript depletion iv.An examination of the impact of BMPR2 transcript depletion on endothelial mitochondrial function by Seahorse assay v.The text of the resubmitted manuscript has also been restructured to ensure that the interpretation of experimental data is more directly in-line with the reported findings.

