## Supplemental Materials for "Circular RNA profiling identifies *circ5078* as a *BMPR2*-derived regulator of endothelial proliferation and stress responses"

### SUPPLEMENTAL MATERIAL

1. Supplemental Methods
2. Supplemental Tables
    - Table S1, Related to Figures 5 and S7
    - Table S2, Related to Figures 1-7 and S1, S2, S5-11
    - Table S3, Related to Figures 1-6, and S2, S5, S8, S10
    - Table S4, Related to Figures 3-7 and S5, S8, S10, S11
    - Table S5, Related to Figures 3-6 and S5, S8, S10
    - Table S6, Related to Figures 1 and S2
3. Supplemental Figures
    - Figure S1, Related to Figure 1
    - Figure S2, Related to Figure 1
    - Figure S3, Related to Figure 2
    - Figure S4, Related to Figure 4
    - Figure S5, Related to Figure 4
    - Figure S6, Related to Figure 5
    - Figure S7, Related to Figure 5
    - Figure S8, Related to Figure 6
    - Figure S9, Related to Figure 6
    - Figure S10, Related to Figure 6
    - Figure S11, Related to Figure 7
4. Supplemental References
5. Supplemental Files
    - File S1, Related to Figures 1-2 and S2
    - File S2, Related to Figures 7 and S11

**Supplemental Methods**

*RNA fluorescence in situ hybridization*

Cells were plated on 12 mm, collagen-coated coverslips (Electron Microscopy Sciences) within the wells of 24 well plates and stained for Caprin-1 protein and linear *BMPR2* RNA using the ViewRNA Cells Plus Assay Kit (ThermoFisher). For the fixation and permeabilization steps, buffers were refreshed halfway through the recommended incubation period. Blocking and incubations with Caprin-1 primary (Proteintech, 15112-1-AP, 1:500 dilution) and Alexa488 conjugated secondary antibodies were performed with buffers containing RNaseOUT RNase inhibitor (both ThermoFisher). Coverslips were mounted with ProLong Diamond Antifade with DAPI (ThermoFisher), and cells were imaged by Stimulated Emission Depletion (STED) super-resolution imaging using a Leica SP8 confocal microscope (Leica Microsystems Inc., ON).

*40S Ribosome Profiling*

Cells were plated and treated with siRNAs and cycloheximide (CHX) as described for polysome profiling. Immediately following CHX treatment, cells were washed once with ice-cold PBS containing 100 μg/mL CHX and fixed on ice in 10 mL of 0.2% paraformaldehyde (PFA, Sigma-Aldrich) for 10 minutes. Fixation was quenched by the addition of glycine to a final concentration of 500 mM, and cells were once again washed with ice-cold PBS containing 100 μg/mL CHX. Cells were then scrape collected in 350 μL of polysome lysis buffer. Lysates were incubated on ice, centrifuged, and RNA content of supernatants determined as described for polysome profiling. 100 μg of total RNA was loaded onto 7.5-30% sucrose gradients (**Table S4**). Gradients were centrifuged at 41,000 rpm for 5 hours at 4°C. Gradients were separated into 800 μL fractions and protein was isolated from these fractions as described for polysome analysis. RNA was precipitated from fractions by overnight incubation with sodium acetate and ethanol at -80°C, before being extracted twice in hot phenol:chloroform:isoamyl alcohol, precipitated in ethanol at -80°C, pelleted, and resuspended in RNase free water[1]. Isolated RNA was reverse transcribed and analyzed by qPCR as described.

### Supplemental Tables

**Table S1: Blood Outgrowth Endothelial Cell (BOEC) Donors**

| Identifier | Group | Age | Sex | Ethnicity | *BMPR2* Mutation | Effect of Mutation |
|---|---|---|---|---|---|---|
| C22 | Control | 22 | F | Caucasian | n/a | n/a |
| C26B | Control | 22 | F | Caucasian | n/a | n/a |
| C33 | Control | 23 | F | Unknown | n/a | n/a |
| C36 | Control | 22 | F | Unknown | n/a | n/a |
| C38 | Control | 47 | F | Caucasian | n/a | n/a |
| C4c | Control | 39 | M | Caucasian | n/a | n/a |
| C50 | Control | 29 | F | Caucasian | n/a | n/a |
| C51 | Control | 29 | M | Caucasian | n/a | n/a |
| C52b | Control | 21 | M | Caucasian | n/a | n/a |
| C54 | Control | 29 | F | Caucasian | n/a | n/a |
| B1 | PAH | 51 | M | Caucasian | R321X | Nonsense |
| B11 | PAH | 32 | M | Caucasian | C.*-944/5GC-AT | Upstream start codon |
| B13 | PAH | 46 | F | Caucasian | E845fsX | Frameshift |
| B4 | PAH | 37 | F | Caucasian | W9X | Nonsense |
| P37-011 | PAH | 72 | M | Caucasian | R321X | Nonsense |

**Table S2: siRNAs and Custom Target Sequences**

| siRNA Name | Catalog Number | Custom siRNA Target sequences |
|---|---|---|
| si*Control* | D-001810-10-20 | n/a |
| si*BMPR2* / si*Linear+5078* | L-005309-00-0005 | n/a |
| si*Linear* | CTM-735230 | GAAACAAGUAGACAUGUAU |
| si*3218* | CTM-713020 | CAACACCACUCACUUCGCAGAAUCA |
| si*5078* | CTM-719614 | AGUAUACAGACAACCUGUCA |
| si*CAPRIN1* | L-016057-00-0005 | n/a |

**Table S3: qPCR Primer Sequences**

| Gene | Forward Primer | Reverse Primer |
|---|---|---|
| *BMPR2* (Linear) | AACATGACAACATTGCCCGC | AAGACGGCAAGAGCTTACCC |
| *BMPR2a* | CAAATCTGTGAGCCCAACAGTCAA | GAGGAAGAATAATCTGGATAAGGACCAAT |
| *BMPR2b* | TGCTGAGGAAAGGATGGCTG | GACTCACCTACGTTCATTCTGC |
| *CAPRIN1* | TGCTACAGAGCAACGACCAC | AGGAAGGGATGATGATGCTG |
| *CCND1* | CCGTCCATGCGGAAGATC | GAAGACCTCCTCCTCGCACT |
| *hsa_circ_0000320* | CATGCCTGATGTGGACCTGA | CAGTCTGGGCCTTGAACCTC |
| *hsa_circ_0000400* | CGACTCTTAGCTTGTCGGGG | CCGTGTTCCAGGCAGTAGAG |
| *hsa_circ_0001218* | GGACATTTTGGTGCCAAGC | TACCTTTCTTGGTCGTGGTG |
| *hsa_circ_0001592* | ATGCGGGTCTTCTTGTTGTC | ATTATGCCGAGCGGGTTG |
| *hsa_circ_0003218* (*circ3218*) | TCCACCTCCTGACACAACAC | CCAAAGGCCATAGCAGGTGC |
| *hsa_circ_0005078* (*circ5078*) | AATGTCCTGGATGGCAGCAG | GGTGTGCTGGACATAGAATGC |
| *hsa_circ_0013876* | AAGATCAAAACCCACCATGC | TGGCCTAAGTCAGGCAGTTC |
| *GAPDH* | AGCCACATCGCTCAGACAC | GCCCAATACGACCAAATCC |
| *RPL19* | AGCGAGCTCTTTCCTTTCG | GAGCCTCTTCTGAAGCCTGA |
| *5.8S rRNA* | ACTCTTAGCGGTGGATCACTC | AAGCGACGCTCAGACAGG |
| *18S rRNA* | CGGCTACCACATCCAAGG | TACAGGGCCTCGAAAGAGTC |
| *28S rRNA* | AGTCGGGTTGCTTGGGAATGC | CCCTTACGGTACTTGTTGACT |

**Table S4: Buffer Compositions**

| Buffer | Components |
|---|---|
| RIPA Buffer | 0.5 M Tris-HCl, pH 8.0 (Sigma-Aldrich)<br>0.1% SDS (Sigma-Aldrich)<br>1% Igepal-CA-630 (Sigma-Aldrich)<br>10 mmol/L sodium fluoride (Sigma-Aldrich)<br>0.5% sodium deoxycholate (Sigma-Aldrich)<br>2 mmol/L sodium orthovanadate (Sigma-Aldrich)<br>1x cOmplete protease inhibitor cocktail (Roche Diagnostics, Basel Switzerland) |
| Homogenization Buffer | 0.25 M Sucrose (Fisher)<br>10 mM Tris-HCl, pH 7.6 (Sigma-Aldrich)<br>100 μg/mL cycloheximide (Sigma-Aldrich)<br>40 U/mL RNasin RNA inhibitor (Promega)<br>1x cOmplete protease inhibitor cocktail (Roche) |
| Cytoskeletal bound polysome (CBP) buffer | 0.25 M Sucrose<br>10 mM Tris-HCl, pH 7.6 (Sigma-Aldrich)<br>130 mM KCl (Sigma-Aldrich)<br>5 mM $MgCl_2$ (Sigma-Aldrich)<br>100 μg/mL cycloheximide (Sigma-Aldrich)<br>40 U/mL RNasin RNA inhibitor (Promega)<br>1x cOmplete protease inhibitor cocktail (Roche) |
| Membrane bound polysome (MBP) buffer | 0.25 M Sucrose<br>10 mM Tris-HCl, pH 7.6 (Sigma-Aldrich)<br>130 mM KCl (Sigma-Aldrich)<br>5 mM $MgCl_2$ (Sigma-Aldrich)<br>0.5% IGEPAL (Sigma-Aldrich)<br>0.5% sodium deoxycholate (Sigma-Aldrich)<br>100 μg/mL cycloheximide (Sigma-Aldrich)<br>40 U/mL RNasin RNA inhibitor (Promega)<br>1x cOmplete protease inhibitor cocktail (Roche) |
| Polysome Lysis Buffer | 20 mM Tris-HCl, pH 7.4 (Sigma-Aldrich)<br>5 mM $MgCl_2$ (Sigma-Aldrich)<br>100 mM KCl (Sigma-Aldrich)<br>1% IGEPAL (Sigma-Aldrich)<br>1 mM Dithiothreitol (DTT)<br>100 μg/mL cycloheximide (Sigma-Aldrich)<br>40 U/mL RNasin RNA inhibitor (Promega)<br>1x cOmplete protease inhibitor cocktail (Roche) |
| 7.5% Sucrose Solution | 20 mM Tris-HCl, pH 7.4 (Sigma-Aldrich)<br>5 mM $MgCl_2$ (Sigma-Aldrich)<br>100 mM KCl (Sigma-Aldrich)<br>7.5% Sucrose (Fisher)<br>100 μg/mL cycloheximide (Sigma-Aldrich)<br>40 U/mL RNasin RNA inhibitor (Promega) |

| | |
|---|---|
| 10% Sucrose Solution | 20 mM Tris-HCl, pH 7.4 (Sigma-Aldrich)<br>5 mM $MgCl_2$ (Sigma-Aldrich)<br>100 mM KCl (Sigma-Aldrich)<br>10% Sucrose (Fisher)<br>100 μg/mL cycloheximide (Sigma-Aldrich)<br>40 U/mL RNasin RNA inhibitor (Promega)<br>1 mM phenylmethylsulfonyl fluoride (PMSF, Sigma-Aldrich) |
| 30% Sucrose Solution | 20 mM Tris-HCl, pH 7.4 (Sigma-Aldrich)<br>5 mM $MgCl_2$ (Sigma-Aldrich)<br>100 mM KCl (Sigma-Aldrich)<br>30% Sucrose (Fisher)<br>100 μg/mL cycloheximide (Sigma-Aldrich)<br>40 U/mL RNasin RNA inhibitor (Promega) |
| 45% Sucrose Solution | 20 mM Tris-HCl, pH 7.4 (Sigma-Aldrich)<br>5 mM $MgCl_2$ (Sigma-Aldrich)<br>100 mM KCl (Sigma-Aldrich)<br>45% Sucrose (Fisher)<br>100 μg/mL cycloheximide (Sigma-Aldrich)<br>4 0U/mL RNasin RNA inhibitor (Promega)<br>1 mM phenylmethylsulfonyl fluoride (PMSF, Sigma-Aldrich) |

**Table S5: Immunoblotting Antibodies**

| Target Antigen | Working dilution | Catalog Number | Supplier |
|---|---|---|---|
| anti-mouse (2°, 700 channel) | 1:10,000 | 926-68070 | LiCOR |
| anti-mouse (2°, 800 channel) | 1:10,000 | 926-32210 | LiCOR |
| anti-rabbit (2°, 700 channel) | 1:10,000 | 925-68071 | LiCOR |
| anti-rabbit (2°, 800 channel) | 1:10,000 | 926-32211 | LiCOR |
| β-actin | 1:1,000 | sc-4778 | Santa Cruz Biotechnology |
| BMPR-II | 1:1,000 | PA521437 | Invitrogen |
| Calnexin | 1:1,000 | 2433 | Cell Signalling Technology |
| Caprin-1 | 1:1,000 | 15112-1-AP | Proteintech |
| Cyclin-D1 | 1:100 | sc-20044 | Santa Cruz Biotechnology |
| eIF2α | 1:1,000 | 9722 | Cell Signalling Technology |
| eIF3B | 1:1,000 | Sc-374156 | Santa Cruz Biotechnology |
| GAPDH | 1:1,000 | 649202 | BioLegend |
| p-eIF2α | 1:500 | 9721 | Cell Signalling Technology |
| RPL7 | 1:2,000 | A300-741A | Bethyl Laboratories |
| RPS6 | 1:1,000 | 2317 | Cell Signalling Technology |

**Table S6: Differential Expression Analysis - circRNAs**

| circRNA annotation | Nearby Gene | baseMean | FoldChange ($log_2$) | lfcSE | stat | *p* - value | $p_{adj}$ |
|---|---|---|---|---|---|---|---|
| *hsa_circ_0000400* | *TUBA1B* | 4446.425847 | -0.500571769 | 0.075126793 | -6.663025897 | 2.68E-11 | 6.72E-09 |
| *hsa_circ_0000320* | *AHNAK* | 1057.53911 | -0.677111846 | 0.101992636 | -6.638830762 | 3.16E-11 | 6.72E-09 |
| *hsa_circ_0005078* | *BMPR2* | 12.33935651 | -5.045507766 | 1.083939309 | -4.654788073 | 3.24E-06 | 0.000459445 |
| *hsa_circ_0013876* | *NBPF10* | 480.5616358 | -0.436167427 | 0.116574816 | -3.74152361 | 0.000182908 | 0.019184544 |
| *hsa_circ_0001218* | *AP1B1* | 132.2569608 | -0.583288546 | 0.158961879 | -3.669361164 | 0.000243157 | 0.019184544 |
| *hsa_circ_0001592* | *H2AC11* | 1185.422388 | -0.346524341 | 0.095154481 | -3.641702825 | 0.000270841 | 0.019184544 |
| *hsa_circ_0001585* | *H2AC6* | 1655.064123 | -0.291130599 | 0.091636627 | -3.177011288 | 0.001488012 | 0.090343612 |
| *hsa_circ_0001242* | *RRP7A* | 343.5215846 | -0.328943783 | 0.108010606 | -3.045476675 | 0.002323118 | 0.123415642 |
| *hsa_circ_0013873* | *NBPF10* | 128.7420834 | -0.47416282 | 0.164850471 | -2.876320689 | 0.004023407 | 0.189994228 |
| *hsa_circ_0000766* | *TUBG1* | 28.89699284 | -0.854375377 | 0.311319724 | -2.744366361 | 0.006062783 | 0.240726774 |
| *hsa_circ_0066631* | *DCBLD2* | 14.69031471 | 1.188562923 | 0.4345126 | 2.735393454 | 0.006230575 | 0.240726774 |
| *hsa_circ_0001243* | *RRP7A* | 449.0279132 | -0.268237588 | 0.099938208 | -2.684034401 | 0.007273962 | 0.257619492 |
| *hsa_circ_0001241* | *RRP7A* | 127.4417904 | -0.460719009 | 0.180527057 | -2.552077331 | 0.010708275 | 0.350078216 |
| *hsa_circ_0072688* | *ADAMTS6* | 182.0977075 | 0.368382891 | 0.148416809 | 2.482083354 | 0.013061672 | 0.382438979 |
| *hsa_circ_0000769* | *TUBG1* | 117.903445 | -0.405071416 | 0.164354288 | -2.464623347 | 0.013715735 | 0.382438979 |
| *hsa_circ_0001591* | *H2AC11* | 63.66976813 | -0.56959946 | 0.232757035 | -2.447184722 | 0.014397703 | 0.382438979 |
| *hsa_circ_0001711* | *VKORC1L1* | 290.0828025 | -0.291525329 | 0.122074806 | -2.388087596 | 0.016936305 | 0.422368499 |
| *hsa_circ_0013872* | *NBPF10* | 525.9883481 | -0.27222644 | 0.114964519 | -2.367917006 | 0.017888548 | 0.422368499 |
| *hsa_circ_0006654* | *SLF2* | 12.73471457 | 1.138444016 | 0.490717063 | 2.319960122 | 0.020343035 | 0.455041565 |
| *hsa_circ_0003692* | *FNDC3B* | 13.76606176 | -1.065342847 | 0.467165694 | -2.280438952 | 0.022581668 | 0.479860437 |
| *hsa_circ_0000767* | *TUBG2* | 731.5344557 | -0.18764512 | 0.085596168 | -2.192214021 | 0.028364057 | 0.556067838 |
| *hsa_circ_0001861* | *GRHPR* | 10.13017975 | 1.192415733 | 0.549177122 | 2.171277145 | 0.029910229 | 0.556067838 |
| *hsa_circ_0002024* | *NAB1* | 10.65052552 | 1.180487012 | 0.54748642 | 2.156194142 | 0.031068509 | 0.556067838 |
| *hsa_circ_0007509* | *PPP4R1* | 12.59314374 | 0.983848703 | 0.457189888 | 2.151947647 | 0.031401478 | 0.556067838 |
| *hsa_circ_0005991* | *APBB2* | 38.63042865 | -0.613831602 | 0.298114242 | -2.059048226 | 0.039489619 | 0.67132352 |
| *hsa_circ_0013878* | *NBPF10* | 1223.221625 | -0.195587561 | 0.097461626 | -2.006816102 | 0.044769248 | 0.715421112 |
| *hsa_circ_0007444* | *RHOBTB3* | 80.31809666 | 0.398395245 | 0.199745021 | 1.994519032 | 0.046095362 | 0.715421112 |

### Supplemental Figures

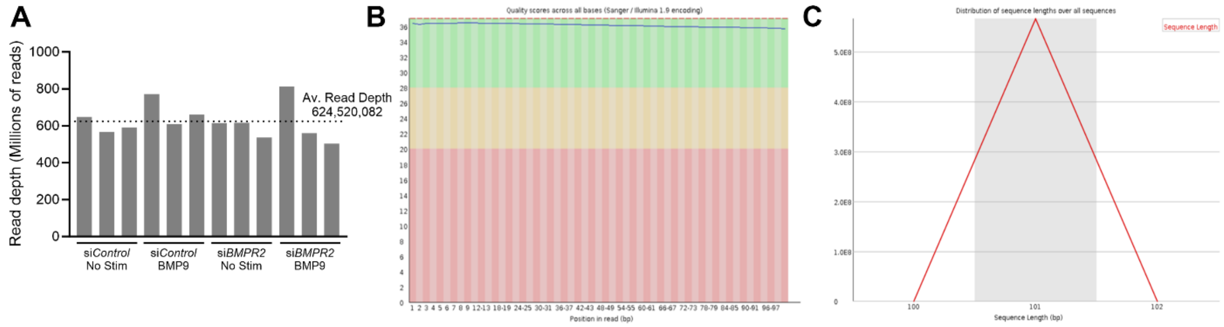


**Figure S1. Assessment of read depth and quality for circRNA screening dataset. (A)** Read depth for each of the 12 samples used in the RNA-seq screen. **(B)** Representative data demonstrating high sequence quality and **(C)** uniform read length, as determined by FastQC.

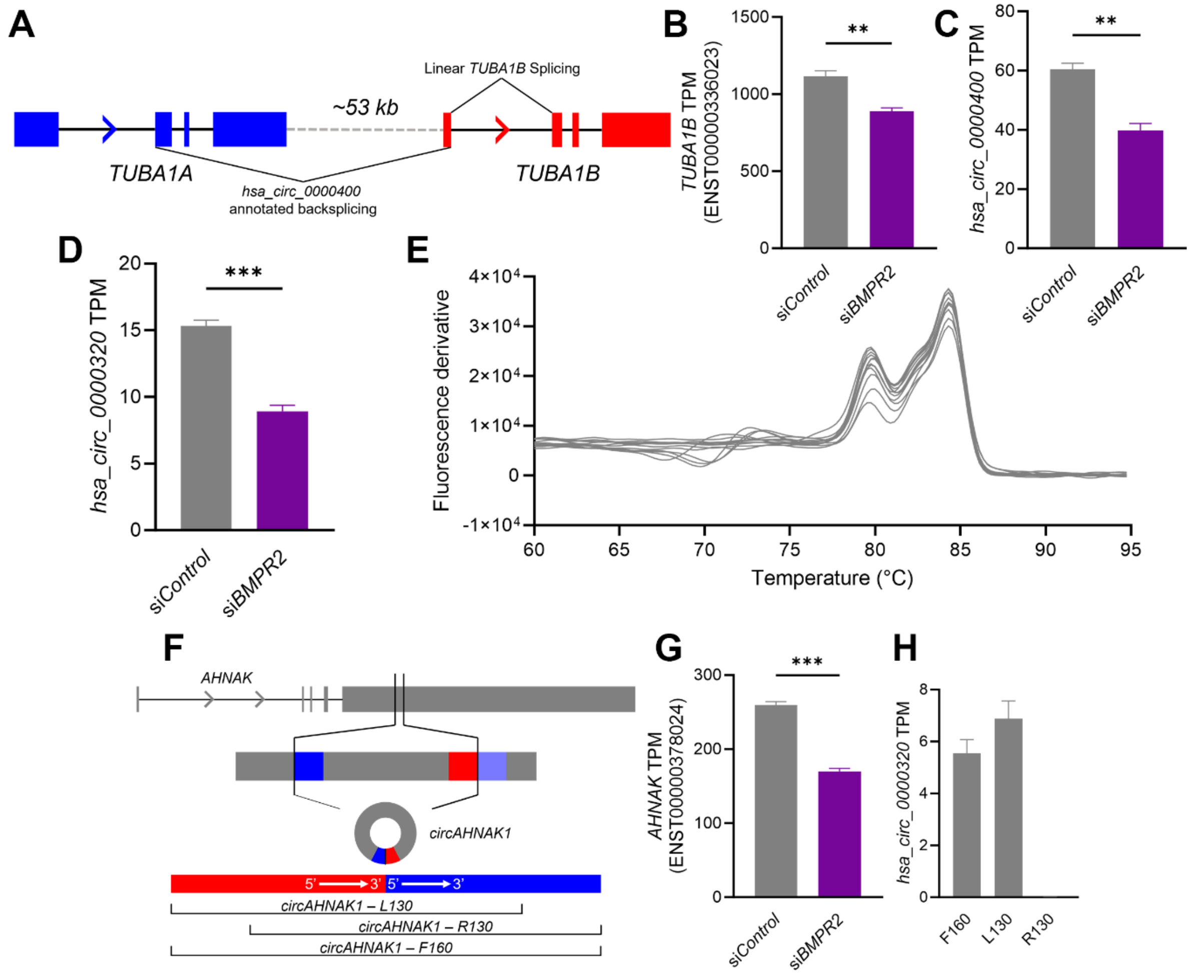


**Figure S2. Examination of differentially expressed circRNA annotations that failed RNase R validation. (A)** The annotation for *hsa_circ_0000400* proposes this circRNA is produced from back splicing between exon 2 of *TUBA1A* and exon 3 of *TUBA1B*, two genes 53kb apart. The proposed back splice junction for *hsa_circ_0000400* produces the same junction sequence as the linear splicing event between exons 1 and 2 of *TUBA1B*. **(B)** Linear *TUBA1B* is dysregulated with *BMPR2* silencing to a similar extent as that of **(C)** the proposed *hsa_circ_0000400* junction (n=3 each). **(D)** The proposed *hsa_circ_0000320* junction is downregulated by *BMPR2* silencing (n=3). **(E)** Melt curve analysis of the PCR product generated using primers reported by Xiao *et. al*. (Aging, 2019) for *hsa_circ_0000320*. Analysis revealed multiple PCR products produced by this primer set (n=6). **(F)** Examination of the *AHNAK* gene from which *hsa_circ_0000320* is derived identified high sequence similarity between the junction sequence produced from the annotated back splicing event (red, blue) and the linear *AHNAK* sequence (red, light blue). **(G)** *AHNAK* was also dysregulated by *BMPR2* silencing, to a similar extent as *hsa_circ_0000320* (n=3). **(H)** The read accumulation for the full 160bp *hsa_circ_0000320* junction sequence (F160) was heavily biased towards the left-most 130bp of the junction (L130) of the junction, which shared an identical sequence with the linear *AHNAK* gene (n=12). **(B-D, H)** Paired t-test. **$p < 0.01$. ***$p < 0.001$.

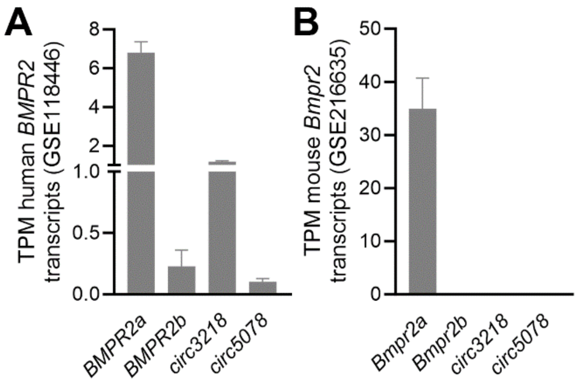


**Figure S3. Linear and circular *BMPR2* RNAs can be detected in other human, but not mouse, endothelial RNA-sequencing datasets.** Transcripts per kilobase million (TPM) of the four *BMPR2*-derived RNAs identified in control HPAECs (n=3) from the data set GSE118446 (Monteiro et. al. *Circ. Res.* 2021), and **(B)** mouse derived endothelial cells (n=4) from the data set GSE216635 (Deng et. al. *Front. Immunol*. 2023).

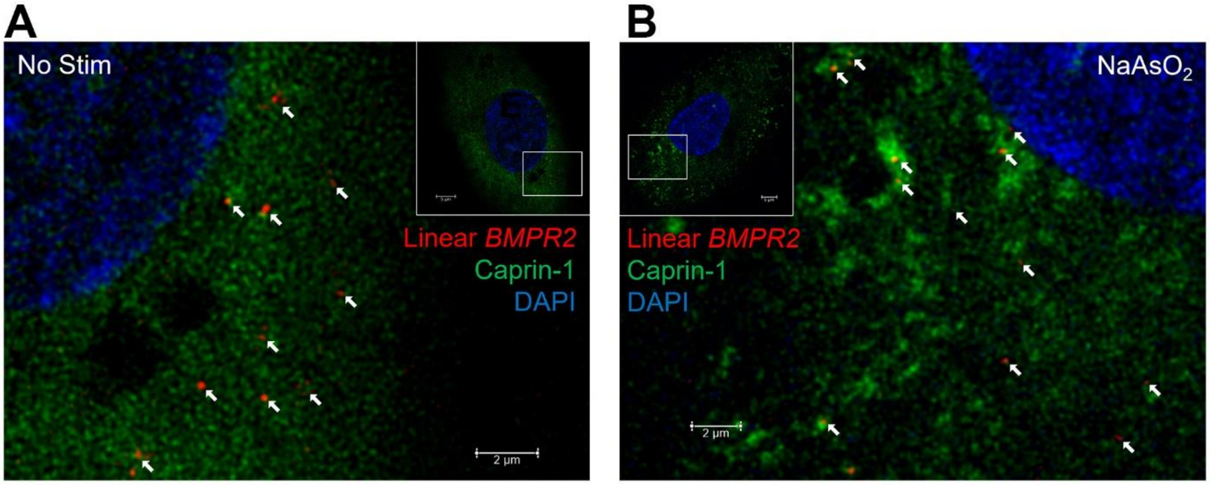


**Figure S4. Fluorescent *in situ* hybridization imaging of Caprin-1 and linear *BMPR2* RNAs. (A)** Stimulated emission depletion (STED) microscopy of untreated HPAECs, with Caprin-1 protein labeled by immunocytochemistry (green) and linear *BMPR2* RNA labeled by fluorescent *in situ* hybridization (red). **(B)** HPAECs labeled in the same manner following 1 hour of treatment with 100 μM sodium arsenite. Inset scale bars are 5 μm. Arrows highlight examples of Caprin-1 co-localization with linear *BMPR2* RNAs.

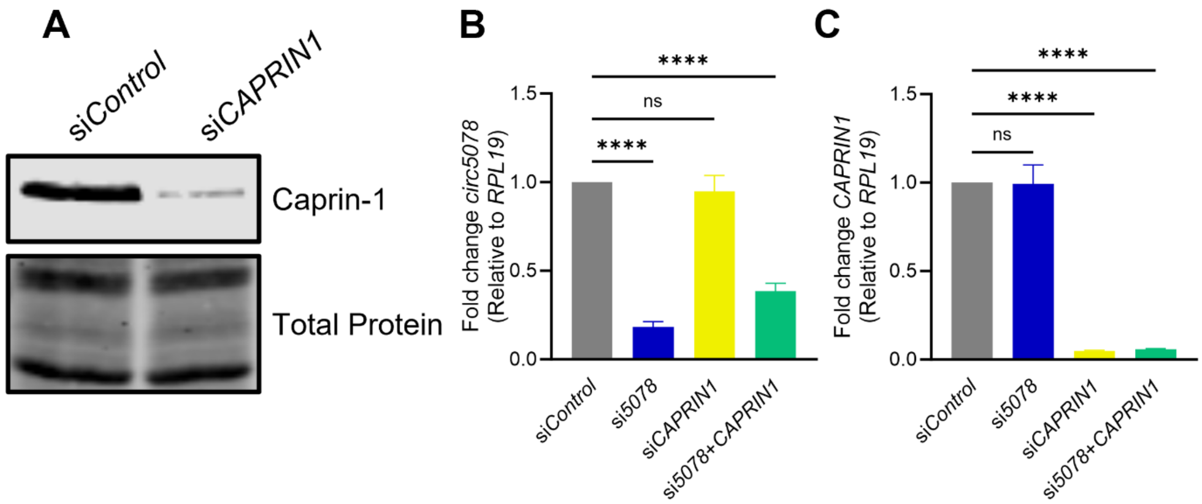


**Figure S5. Validation of Caprin-1 antibody and *circ5078*/*CAPRIN1* double knockdowns. (A)** Immunoblot of Caprin-1 in HPAECs treated with si*Control* or si*CAPRIN1*. **(B)** Quantification of *circ5078* expression and **(C)** *CAPRIN1* expression in HPAECs treated with siRNAs targeting *circ5078*, *CAPRIN1*, or both (n=8). **(B, C)** one-way ANOVA with Dunnett's post hoc test. ns indicates not significant. ****$p < 0.0001$.

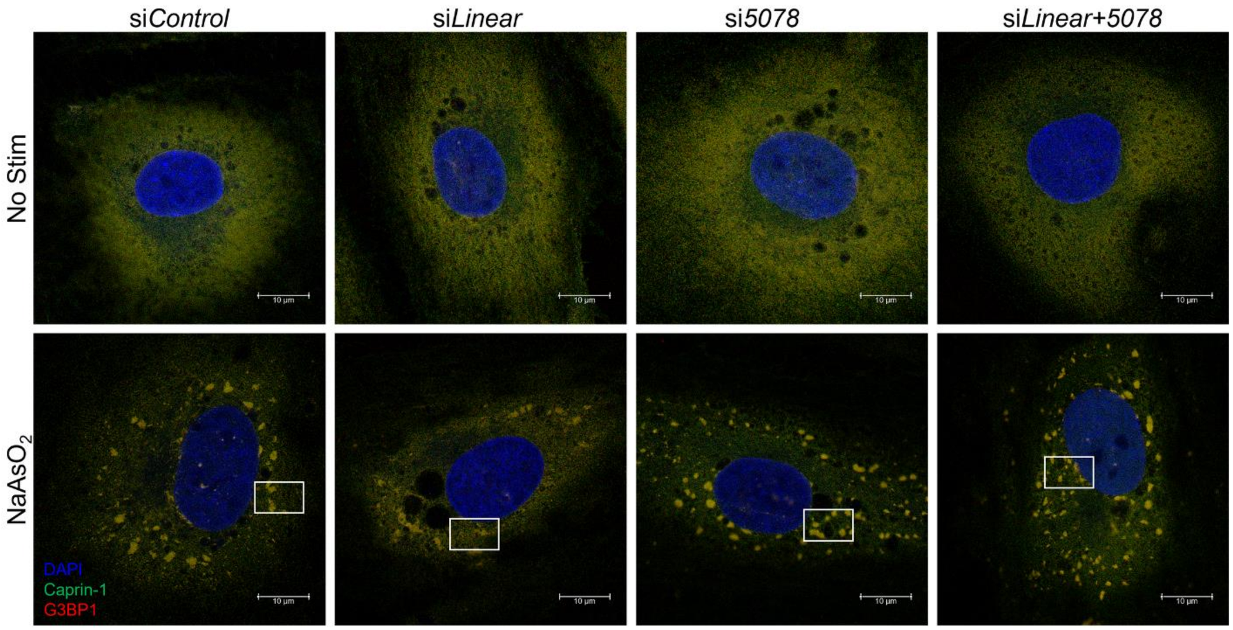


**Figure S6. Silencing of *BMPR2* transcripts influences the formation of cytoplasmic stress granules in response to sodium arsenite treatment.** Representative stimulated emission depletion (STED) microscopy images of single cells under si*Control*, si*Linear*, si*5078* or si*Linear+5078* silencing conditions, with and without sodium arsenite treatment. Highlighted boxes indicate the areas from which the images in **Figure 4E** are taken.

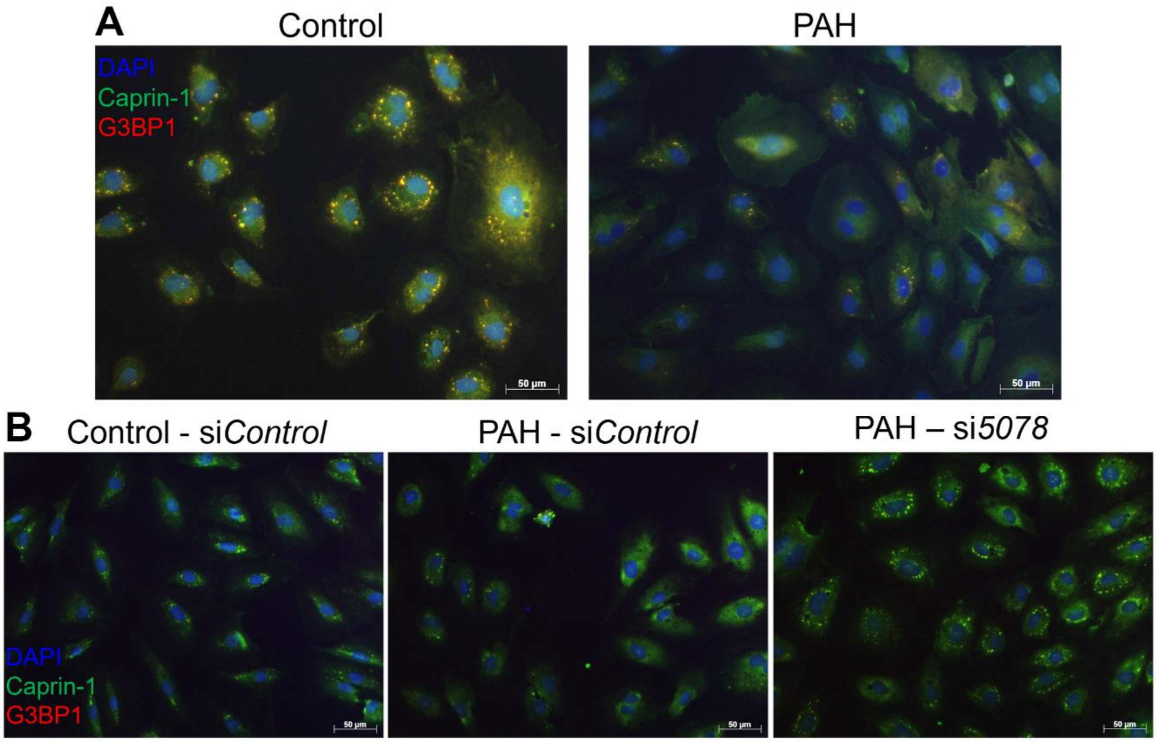


**Figure S7. The relative abundance of linear *BMPR2* and *circ5078* regulate the endothelial stress response in PAH. (A)** Representative epifluorescence images of blood outgrowth endothelial cells (BOECs) from healthy controls and PAH patients following treatment with sodium arsenite. **(B)** Representative images of control and PAH derived BOECs following siRNA and sodium arsenite treatment.

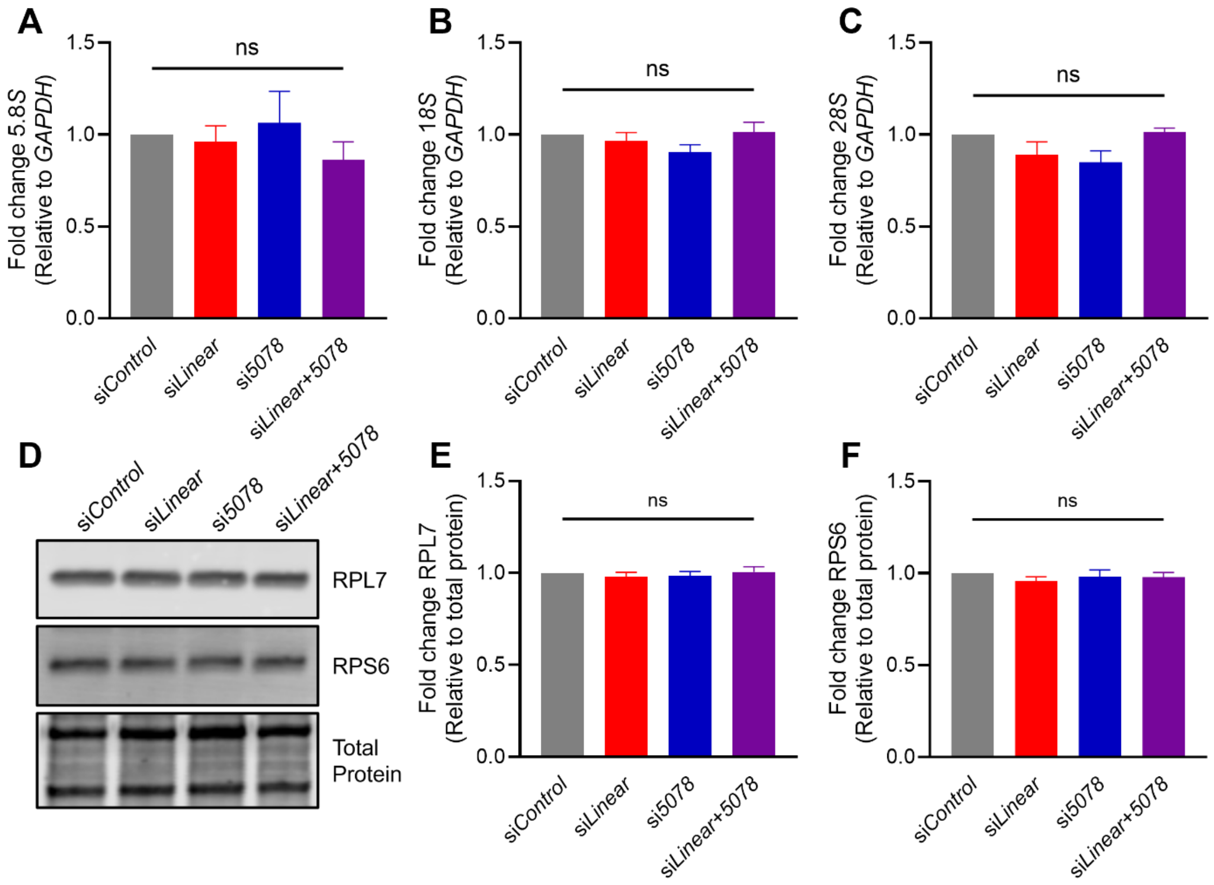


**Figure S8. Ribosome biogenesis is not impacted by the loss of *BMPR2*-derived transcripts.** **(A)** Relative expression of *5.8S*, **(B)** *18S*, and **(C)** *28S* ribosomal RNAs in HPAECs following the silencing of linear *BMPR2* transcripts, *circ5078* or both, compared to si*Control* treated cells (n=6). **(D)** Representative immunoblot and quantification of **(E)** RPL7 and **(F)** RPS6 protein expression under these silencing conditions (n=8). **(A-C, E, F)** One-way ANOVA with Dunnett's post-hoc test. ns indicates not significant.

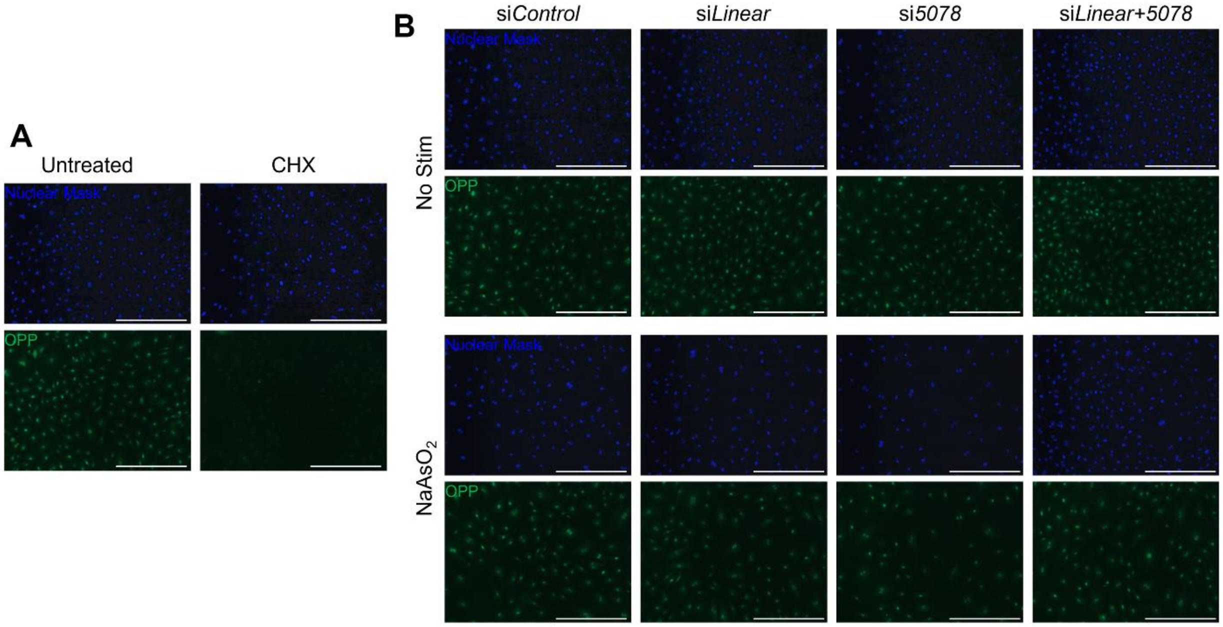


**Figure S9. Loss of *BMPR2*-derived RNAs does not alter global protein synthesis. (A)** Representative epifluorescence images of HPAECs stained with Nuclear Mask and Alexa488 labeled O-propargyl-puromycin (OPP). Cells were left untreated or subjected to 100 µg/mL cycloheximide (CHX) for 1 hour, with OPP added for the final 30 minutes. **(B)** Representative images of HPAECs treated with *BMPR2*-targeting siRNAs, with and without 1 hour of 100 µM of sodium arsenite. OPP incorporation was quantified for the final 30 minutes of this treatment, as in (**A**). Scale bars are 400 µm.

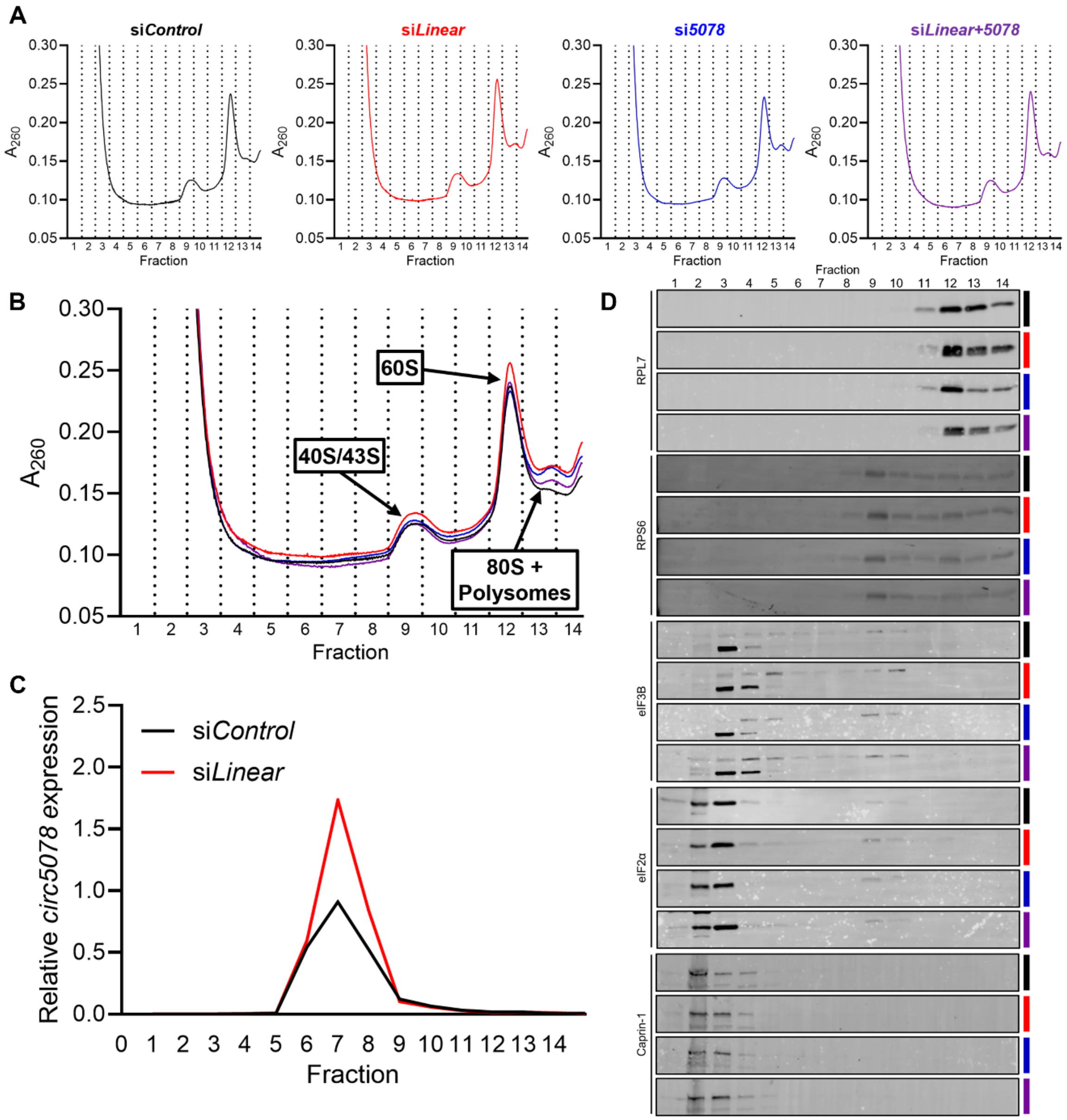


**Figure S10. *circ5078* accumulates in non-ribosomal complexes. (A)** Representative UV absorbance ($A_{260}$) profiles from 7.5-30% sucrose gradients of HPAECs treated with si*Control* (black), si*Linear* (red), si*5078* (blue), and si*Linear+5078* (purple). **(B)** Overlay of $A_{260}$ profiles highlighting the 43/48S preinitiation complex (PIC), 60S subunits, and 80S monosomes and polysomes. **(C)** Distribution of *circ5078* across 7.5-30% sucrose gradients in HPAECs treated with si*Control* or si*Linear*. Plot is the average of three independent experiments. **(D)** Representative immunoblots of RPL7, RPS6, eIF3B, eIF2α and Caprin-1 across each fraction under each knockdown condition

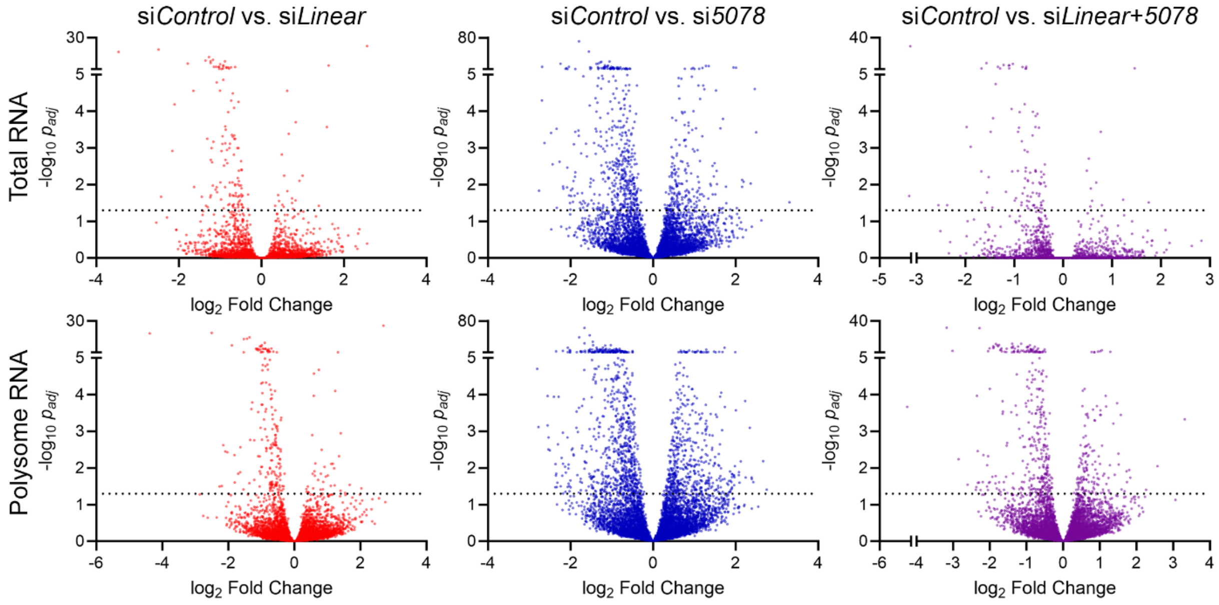


**Figure S11. *BMPR2* transcript depletion induces changes to total cellular mRNA expression and the translational efficiency of select mRNAs.** Volcano plots of total endothelial mRNA (top) and mRNA from actively translated polysome fractions (Fractions 8-13, bottom) in response to silencing of linear *BMPR2* (left), *circ5078* (middle), or both linear *BMPR2* and *circ5078* (right). Dashed line represents $p_{adj}$ = 0.05.
